# Systemic RNAi in planarians depends on spread of RNPs from active stem cells

**DOI:** 10.64898/2026.03.03.709314

**Authors:** Sudheesh Allikka Parambil, Karina Ascunce Gonzalez, Axel Poulet, Levi Cruz, Hae-Lim Lee, Mariana Witmer, Josien C. van Wolfswinkel

**Affiliations:** Department of Molecular Cellular and Developmental Biology, Yale University, New Haven CT 06511, USA; Yale Stem Cell Center, Yale School of Medicine, New Haven CT 06511, USA; Yale Center for RNA Science and Medicine, Yale School of Medicine, New Haven CT 06511, USA

## Abstract

RNAi is a powerful cellular defense mechanism against genomic invaders that rely on dsRNA intermediates, such as viruses and mobile elements. Due to its specificity and ease of use, RNAi is also widely used as an experimental and therapeutic strategy to reduce levels of specific RNAs. For many systems however, delivery of the silencing agents into each individual cell is a major challenge. A mechanistic understanding of the processes involved in intercellular spread in systems with effective systemic RNAi thus is key to improve applications, and to increase our understanding of this defense mechanism. Remarkably, outside of *C. elegans* and plants, our molecular understanding of systemic RNAi remains very limited.

We here investigated the highly-effective systemic RNAi of the planarian *S. mediterranea*, which can be triggered by introduction of dsRNA via food or injection and rapidly spreads through the entire body. Notwithstanding its efficiency, we find no evidence of an RNA amplification mechanism or of transgenerational effects as are found in *C. elegans*, and rather find that planarian RNAi effects are limited in time. We identify the biogenesis factors involved in the planarian RNAi mechanism, and find that these are independent of the miRNA pathway, enabling the separation of the effects from these small RNA pathways. Surprisingly, we find that planarian systemic RNAi relies on active stem cells. Further, we identify Argonaute-siRNA complexes as the mobile agent that effectuates systemic spread of RNAi throughout the tissue.

These findings provide new insights into the mechanisms by which small RNAs spread between cells, and by which organisms can extend protection to all their cells upon encounter of a novel invading element. Additionally, our findings may have important implications for the design of effective applications in other systems.

## Introduction

Small RNA mediated gene regulation is an essential mechanism in metazoan development. It forms a crucial layer of posttranscriptional gene control that finetunes and checks the outcomes of the transcriptional regulatory processes. This allows cells to eliminate detrimental transcripts, rapidly adapt to changing conditions, and to dampen gene expression noise, resulting in cleaner developmental transitions and more robust development (Tam et al. 2008; Schmiedel et al. 2015). Additionally, small non-coding RNAs function in the defense against invading genomic elements (Girard and Hannon 2008; Malone and Hannon 2009; van Wolfswinkel and Ketting 2010). They can dynamically recognize and neutralize such elements by targeting sequence motifs found in invading elements and thereby form an essential nucleic-acid centered immune system.

Small non-coding RNAs exist in three main flavors: microRNAs (miRNAs), small interfering RNAs (siRNAs), and PIWI interacting RNAs (piRNAs) (Ghildiyal and Zamore 2009; Kim et al. 2009). piRNAs are primarily present in the animal germline and their biogenesis machinery is distinct from that of the other two types of small RNAs. miRNAs and siRNAs however are similar in many regards. The core of these silencing mechanisms consists of two factors: an RNA endonuclease named Dicer that cuts fragments of around 22nt from a larger piece of dsRNA or hairpin RNA; and an RNA-binding protein of the Argonaute family that binds one of the small RNA strands and uses this to identify target sequences by base-pairing (Hutvagner and Simard 2008; Carthew and Sontheimer 2009). miRNAs and siRNAs however differ in the origin of their Dicer substrate and in their molecular outcome. Whereas miRNAs are encoded in the genome and function primarily in cell differentiation and developmental transitions, siRNAs are derived from any endogenous or exogenous sources of double stranded RNA (dsRNA). dsRNA is commonly produced as a consequence of transposon activity or viral replication, and siRNAs thus function primarily in the defense against mobile elements and viruses (Chen and Hur 2022). Once bound by an Argonaute protein, they identify target RNAs by perfect base-pairing, resulting in cleavage and degradation of the target RNA and the siRNA-loaded Argonaute is released to cleave another target molecule.

In plants and nematodes, the Argonaute recognition of a target RNA triggers an amplification loop that appears to be absent from most metazoans, including humans. This amplification mechanism involves an RNA dependent RNA polymerase (RdRP) that generates new RNA strands complementary to target RNA molecules, thereby increasing the amount of antisense siRNA, while also expanding the targeted region outside the limits of the original dsRNA trigger (Dalmay et al. 2000; Sijen et al. 2001; Shukla et al. 2020). This amplification mechanism makes RNAi highly effective and allows for powerful systemic and transgenerational effects triggered by dsRNA in *C. elegans* and in plants. Whether the presence of an amplification mechanisms is a requirement for highly effective RNAi-mediated silencing however remains unclear, as a detailed understanding of the RNAi mechanism in other systems is lacking.

An important aspect of small RNA-mediated silencing is that it can spread between cells, which may be particularly important to halt the propagation of viruses or activated mobile elements.

Remarkably, the mechanism of systemic RNAi remains poorly understood. The nature of the migrating molecules, as well as the mechanisms by which they are secreted and taken up remain enigmatic. The process has been best studied in *C. elegans* where dsRNA is imported from the extracellular space with the help of the transmembrane protein SID-1 (Winston et al. 2002).

There are however indications that other mobile entities which have remained unresolved take part in the systemic silencing (Jose et al. 2011). In plants, small RNAs are thought to spread directly between cells through plasmodesmata, but longer distance migration through the phloem has also been described (Molnar et al. 2010). Whether these migrating small RNAs are part of any type of complex, and how they are released or imported remains unclear. Improved knowledge of the mechanisms of small RNA mobility and uptake is important for our understanding of viral defense mechanisms, and additionally can provide valuable insights for delivery strategies of therapeutic applications using small RNAs.

In the planarian *S. mediterranea*, the RNAi pathway is highly effective for gene knockdown, and is routinely used for forward genetics (Sanchez Alvarado and Newmark 1999; Newmark et al. 2003). dsRNA introduced by food or injection efficiently induces silencing, and is thought to reach every cell of the body. However, the identity of the silencing agent that spreads between the cells has remained unknown. Further, the high efficiency of the knockdown effects has led to the notion that an amplification mechanism must be present in planarians, even though no RdRP homologs can be identified by sequence similarity (Fontenla et al. 2017; Pinzon et al. 2019).

We here used the planarian *S. mediterranea* to investigate the mechanisms involved in effective systemic RNAi in the absence of an RdRP. We find that *S. mediterranea* uses distinct Argonaute proteins and distinct Dicers for the biogenesis of miRNAs and siRNAs, facilitating the separation of these two pathways. We find that initiation of RNAi requires the dsRNA transporter SID-1 and vesicular transport, as well as the Argonaute proteins AGO-1 and AGO-3, and the Dicer protein DCR-2. Remarkably, we find that actively cycling stem cells are essential for systemic RNAi. We additionally find that systemic RNAi involves the transport of loaded Argonaute complexes between cells and no longer requires SID-1 or Dicer in the recipient cells. The RNPs that transfer silencing are sensitive to RNAse and protease, indicating that membrane encapsulation is not required for their transport or uptake.

Our findings give new insight in the functioning of RNAi and provide indications for the endogenous function of the RNAi mechanism in planarians. Moreover, they reveal the workings of a highly effective siRNA distribution system, which may have important implications for the design of effective RNAi-based therapies.

## Results

### The planarian siRNA and miRNA pathways use separate molecular machinery

The main effector proteins in the RNAi pathway are the Dicer protein, which cuts the dsRNA into ∼22nt fragments, and the Argonaute protein which binds the ∼22nt siRNA and uses it to identify a target for silencing. The same molecular functions however are required in the miRNA pathway, and in many organisms these two pathways share the use of the same proteins, making it difficult to separate their effects (**Figure 1A**).

**Figure 1.**
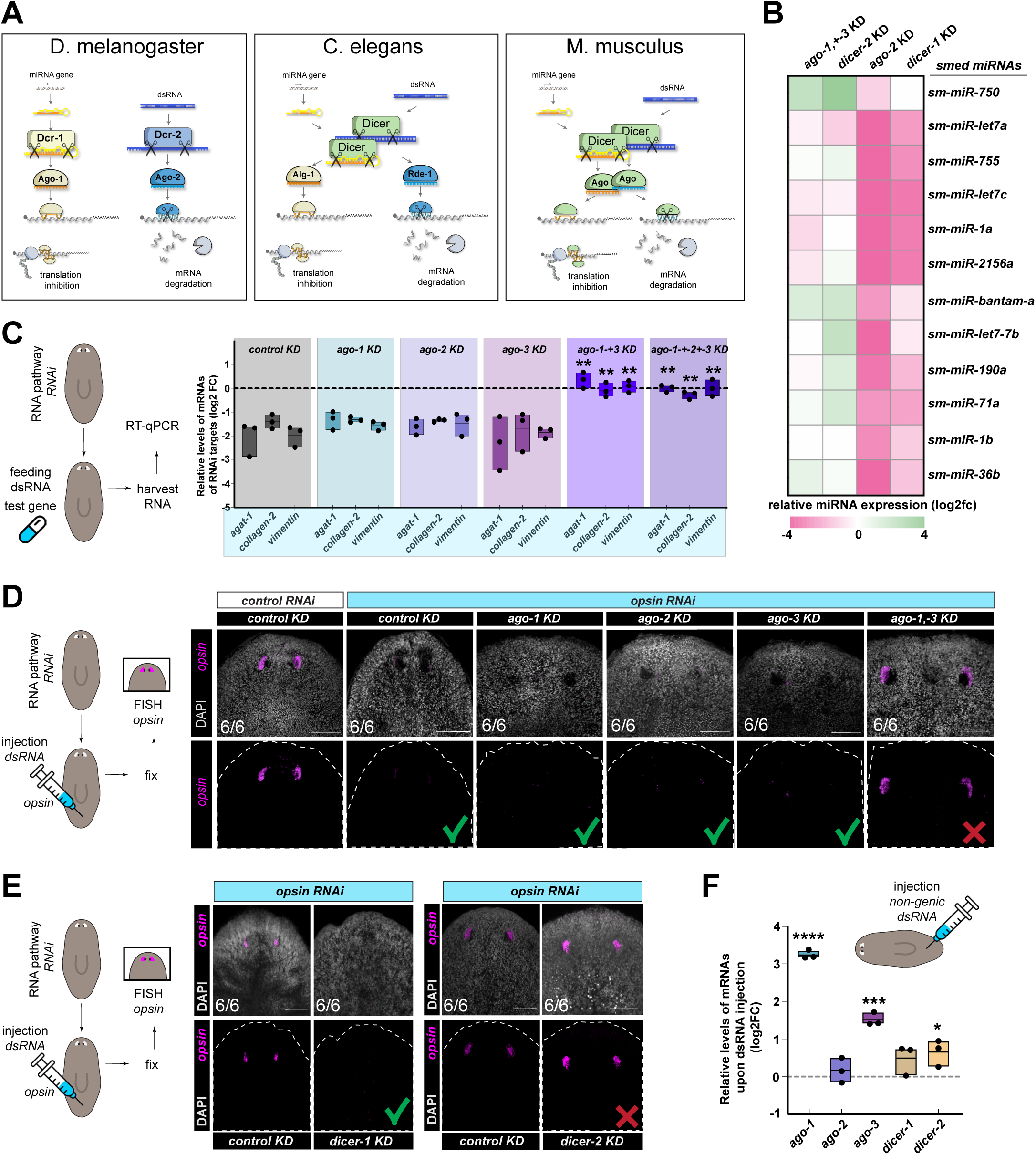
Planarian siRNA and miRNA pathways use separate molecular machinery. **A.** Schematic showing the overlap in involvement of Dicers and Argonaute proteins between the miRNA and siRNA pathways in several model systems. **B.** Heatmap showing the log2 fold changes in miRNA levels as determined by qPCR upon knockdown of *ago-1+3*, *ago-2*, *dcr-1*, or *dcr-2*, relative to control samples. See also Suppl Figure S2A. **C.** Boxplots showing the efficiency of RNAi-mediated knockdown of test transcripts *agat-1*, *collagen-2*, and *vimentin-1*, in control animals, or upon knockdown of *ago-1*, *ago-2*, *ago-3*, *ago-1+3*, or *ago-1+2+3*, as determined by qPCR. Absence of an individual Argonaute does not affect the efficiency of RNAi, but elimination of AGO-1 and AGO-3 simultaneously results in ineffective RNAi. Significance: **p ≤ 0.01 by Student’s t-test, comparison between control and *ago* knockdown. **D.** Experimental schematic (left), and confocal FISH images (right) showing the efficiency of RNAi-mediated knockdown of eye transcript *opsin*, in control animals, or upon knockdown of *ago-1*, *ago-2*, *ago-3*, or *ago-1+3*. Absence of an individual Argonaute does not affect the efficiency of RNAi, but elimination of AGO-1 and AGO-3 simultaneously results in ineffective RNAi. Green check marks indicate effective RNAi; red crosses indicate ineffective RNAi. **E.** Experimental schematic (left), and confocal FISH images (right) showing the efficiency of RNAi-mediated knockdown of eye transcript *opsin*, in control animals, or upon knockdown of *dcr-1* or *dcr-2*. Absence of DCR-1 does not affect the efficiency of RNAi, but elimination of DCR-2 results in ineffective RNAi. Green check marks indicate effective RNAi; red crosses indicate ineffective RNAi. **F.** Box plot showing changes in expression levels of Argonaute and Dicer genes (log2 fold changes) upon injection of unrelated dsRNA into the animals. Significance: *p ≤ 0.05, **p ≤ 0.01, ***p ≤ 0.001, ****p ≤ 0.0001, by Student’s t-test.

Inspection of the planarian genome identified three Argonaute proteins, and two Dicers (**Suppl Figure S1**). In phylogenetic analysis, one of the Argonaute proteins clustered with the vertebrate Argonautes as well as the miRNA-binding Argonautes of other invertebrate models. This Argonaute was previously named *ago-2* (Li et al. 2011) and was proposed to function in stem cell maintenance. The other two Argonautes (*ago-1* and *ago-3*) were found adjacent to each other in the genome, and in phylogenetic analysis clustered together, away from the miRNA-binding Argonautes and closer to the nematode-specific WAGOs (**Suppl Figure S1A**). All three Argonautes contained the catalytic DDH triad, suggesting that they could function in target cleavage. Structural predictions revealed that, as expected by the phylogenetic analysis, AGO-2 closely resembled vertebrate Argonautes, whereas AGO-1 and AGO-3 had more divergent PAZ and MID domains (**Suppl Figure S1B,C**).

To determine whether any of the planarian Argonaute proteins were specific to the miRNA or the siRNA pathway, we first used qPCR to measure changes in the levels of a set of miRNAs upon Argonaute knockdown (**Suppl Figure 2A-C**). We found that knockdown of *ago-1, ago-3,* or *ago-1* and *ago-3* combined did not induce major changes in miRNA levels, nor did it produce any observable phenotypes even after 60 days of RNAi (**Figure 1B, Suppl Figure 2B-D).** In contrast, *ago-2(RNAi)* animals began showing head regression around day 28 of RNAi treatment, eventually resulting in lethality (**Suppl Figure 2B,C**). Additionally, knockdown of *ago-2* resulted in a decreased level of each of the miRNAs tested, indicating that planarian miRNAs rely primarily on AGO-2 (**Figure 1B, Suppl Figure 2D**)(Li et al. 2011).

We next tested the effect of Argonaute knockdown on the efficiency of RNAi (**Figure 1C, Suppl Figure S2E,F**). We used qPCR to measure the effect of loss of any of the Argonautes on the knockdown efficiency of the epidermal genes *agat-1* and *vimentin-1*, and the muscle gene *collagen-2*. We found that animals deficient for any of the individual Argonautes remained proficient in RNAi, but animals deficient in *ago-1* and *ago-3* simultaneously were insensitive to subsequent dsRNA of the test genes (**Figure 1C**). As a complementary strategy we determined the efficiency of knockdown of the eye gene *opsin* as analyzed by FISH (**Figure 1D**), and similarly found that RNAi was ineffective in the *ago-1&-3* simultaneous knockdown. These results indicate that AGO-2 is not involved in RNAi, but AGO-1 and AGO-3 both function in the RNAi pathway and are (partially) redundant.

*S. mediterranea* encodes two Dicers (in addition to the related protein Drosha that functions specifically in the miRNA pathway). In phylogenetic analysis, one of the Dicers (*dcr-1*) clustered with the vertebrate Dicers, whereas the other (*dcr-2*) was more divergent (**Suppl Figure S1D-F**). We wondered whether, similar to the Argonautes, the Dicers might each function in only one of the small RNA pathways. Indeed, we found that most miRNAs were reduced in level upon knockdown of *dcr-1*, but not upon knockdown of *dcr-2* (**Figure 1B, Suppl Figure S2G-I**), whereas knockdown of *dcr-2* (but not *dcr-1*) resulted in insensitivity to other dsRNA triggers (**Figure 1E**).

Remarkably, we found that the factors that we had identified in as part of the siRNA pathway (*ago-1*, *ago-3*, and to a lesser extent *dcr-2*), also increased in expression level upon injection of dsRNA, whereas the miRNA-related factors *ago-2* and *dcr-1*) showed no significant response (**Figure 1F**). This suggests a feedback loop where increased presence of dsRNA results in increased production of the proteins involved in its processing and defense.

Together, these analyses show that the miRNA pathway and the siRNA pathway in *S. mediterranea* use distinct Argonautes and Dicers. This thus allows us to study the mechanism of RNAi in planarians without abolishing miRNA function.

### Planarian RNAi is durable and systemic, but typically not heritable

Due to the apparent efficiency of planarian RNAi (Sanchez Alvarado and Newmark 1999; Newmark et al. 2003), it has often been posed that *S. mediterranea* must employ a mechanism to amplify the silencing. However, since the original papers describing the effectivity of RNAi, mechanistic insights have remained sparse. We therefore first set out to establish how far the effects of a dsRNA trigger really extend. For ease of detection we focused on genes expressed in the planarian eyes that can be visualized by fluorescent in situ hybridization (FISH) (**Figure 2A, Suppl Figure S3A,B**) (Lapan and Reddien 2012), and for consistency, we used injection into the body cavity as the method to introduce the dsRNA. Consistent with previous reports, we found that RNAi of the eye-specific gene *opsin* resulted in loss of transcript from the eyes within 16 hours (**Figure 2B, Suppl Figure S3B**) (Sanchez Alvarado and Newmark 1999; Rouhana et al. 2013). The targeted mRNA remained suppressed until at least 12 days later, but by 16 days post injection, the mRNA could be detected again. Similar dynamics were detected using the pharynx gene laminin as the RNAi target (**Suppl Figure S3C**). Above a threshold of ∼10^10^ molecules per animal dsRNA concentration did not significantly affect the dynamics (**Suppl Figure S3D**). If the animals went through regeneration after injection of dsRNA, the silencing effect was extended to the newly formed tissue, but the time window of the RNAi efficacy was not significantly altered (**Figure 2C, Suppl Figure S3E**). Altogether, this suggests that the temporal window of operation for RNAi in planarians is quite consistent and is around two weeks, independent of tissue or dsRNA concentration.

**Figure 2.**
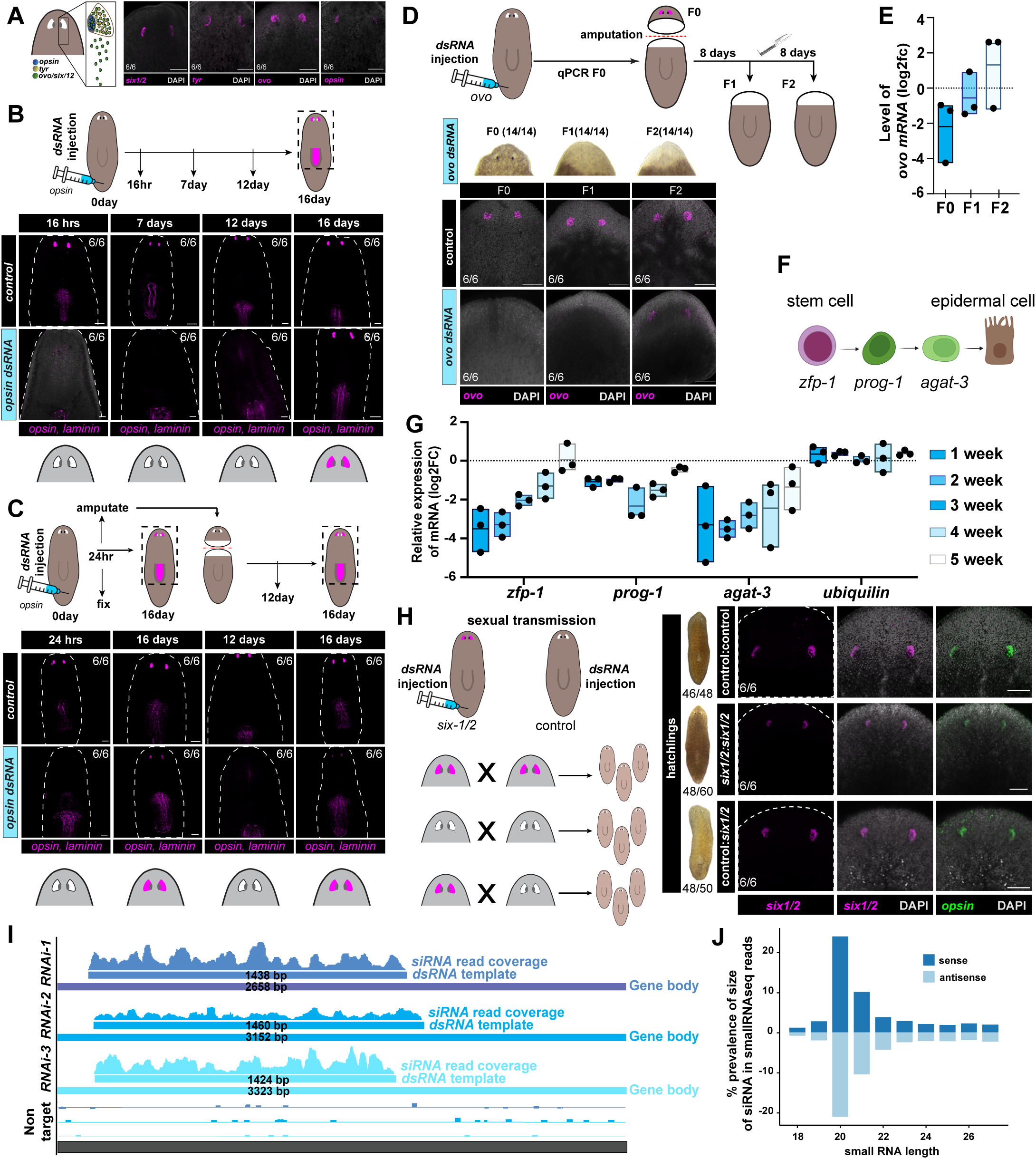
Planarian RNAi is durable and systemic, but typically not heritable **A.** Experimental schematic (left), and confocal FISH images (right) showing the expression patterns of eye genes used in this study. RNAs for transcription factors *ovo* and *six-1/2* label eye progenitors, whereas *opsin* and *tyronsinase hydroxylase* (*tyr*) mark differentiated eye cells. **B.** Experimental schematic (top), and confocal FISH images (bottom) showing expression level of *opsin* at different times upon injection of *opsin* dsRNA. The expression of non-target control laminin remained unchanged, demonstrating RNAi specificity. **C.** Experimental schematic (top), and confocal FISH images (bottom) showing expression level of *opsin* at different times upon injection of *opsin* dsRNA in homeostatic animals or in animals subjected to amputation indicate that the duration of RNAi-mediated knockdown is not altered by regeneration. **D.** Experimental schematic (top) illustrating the paradigm to show downstream effects of RNAi-mediated knockdown of eye progenitors. Serial amputation products are labeled F1 and F2. Brightfield images (middle) show macroscopic eyes, and confocal FISH images (bottom) show expression levels of *ovo* upon injection of *ovo* dsRNA followed by repeated head amputation and regeneration. Eyes remain affected by the RNAi-mediated knockdown in the F2, but *ovo* mRNA has returned at that time. **E.** Box plot showing the change in *ovo* RNA level (log2 fold change relative to t=0) upon injection of *ovo* dsRNA followed by repeated cycles of head amputation and regeneration (F1-F2). While macroscopic eyes remain affected, *ovo* mRNA levels recover after one round of regeneration. **F.** Schematic showing stages of epidermal cell differentiation and characteristic mRNAs. **G.** Box plot showing the change in RNA levels (log2 fold change relative to t=0) of control gene *ubiquilin* and epidermal linage genes *zfp-1*, *prog-1*, and *agat-3* over the course of 5 weeks after feeding of *zfp* dsRNA. **H.** Experimental schematic (left), and confocal FISH images (right) showing expression level of *six-1/2* and *opsin* in the progeny of animals subjected to injection of *six-1/2* dsRNA or control dsRNA. **I.** Normalized genome browser views of 3 different RNAi experiments, showing the targets and the mapping of small RNAs matching these targets in small RNA libraries from the respective RNAi experiment. **J.** Bar graph showing the size and orientation (sense above the axis; antisense below the axis) of the small RNA reads matching the target in aggregated RNAi experiments.

Although direct effects of dsRNA lasted for only two weeks, downstream effects could be detected over longer timespans. Injection of dsRNA targeting the transcription factor *ovo*, which is required for the differentiation of eye progenitors, did not remove existing eyes in homeostatic animals, but did prohibit the formation of eyes in newly regenerating heads (**Figure 2D, Suppl Figure S4A)**. These heads then remained eye-less for extended amounts of time. This phenotype was reproduced using another eye-specific transcription factor as the dsRNA target (*six-1/2*, **Suppl Figure S4B,C**), but not by targeting non-transcription factor genes specific to the eyes (*opsin*, *tyr* **Figure 2C**). Repeated amputations could result in repeated absence of the eyes in newly regenerating heads, but, as expected based on the previously established duration of RNAi effects, if the amputation took place at 16 days after the initial RNAi injection normal eye were formed (**Suppl Figure S4B**). Additionally, we found that in the regenerating eye-less animals, the mRNA level of the transcription factor itself was restored to homeostatic levels by 16 days after injection (**Figure 2D,E, Suppl Figure S4D**), indicating that the sustained absence of macroscopic eyes in regenerating heads was not a reflection of extended RNAi, but rather was a downstream developmental phenotype.

If dsRNA was introduced by feeding, the silencing effects lasted notably longer. When *opsin* dsRNA was delivered by feeding, the silencing effects lasted around 3 weeks instead of the 2 weeks found by injection (**Suppl Figure S4E**). Reduction of the neoblast gene *zfp-1* could still be detected at 4 weeks after the last dsRNA feeding (**Figure 2F,G**). By 5 weeks, the expression level of the targeted mRNA was back to homeostatic levels, but effects on downstream genes that were regulated by the targeted gene, could again last well beyond this timeframe. For example, we found that the level of the epidermal progenitor marker *agat-3* (**Figure 2F**) remained detectably reduced at 5 weeks after the end of dsRNA feedings of the epidermal master regulator *zfp-1*, even though the level of *zfp-1* itself was no longer reduced (**Figure 2G**).

As transmission of RNAi-mediated silencing was detected through regeneration, which is the asexual mode of propagation in planarians, we wondered whether RNAi could be transmitted through sexual generations. To test this, we switched to the related species *Schmidtea polychroa* which is more fertile. We established that RNAi of the transcription factor *six-1/2* resulted in a strong reduction of the *six-1/2* mRNA (**Suppl Figure S4F**), and in the absence of newly formed eyes in regenerating heads (**Suppl Figure S4G,H**), as it did in *S. mediterranea*. If such animals were crossed with other *six-1/2(RNAi)* or control animals however, most of the resulting hatchlings had normal eyes that were indistinguishable from eyes of control hatchlings, indicating that the RNAi effect was not broadly transmitted to the next generation (**Figure 2H**). The only exception were hatchlings from capsules laid within 2 days after feeding. Several of these hatchlings did not develop normal eyes, suggesting that some transmission is possible during the time that dsRNA levels in the parent are high. It is possible that such dsRNA is loaded into the capsules and continues to act on the developing hatchling.

The effectiveness of RNAi-mediated silencing in *S. mediterranea* has raised the suggestion that an amplification mechanism for small RNAs, similar to the situation in *C. elegans* or plants might be in place. To evaluate evidence for such an amplification mechanism, we inspected the siRNA reads in small RNA sequencing data from RNAi experiments. If amplification of small RNA were involved, we would expect such small RNAs (1) to have an antisense strand bias and (2) to not be strictly limited to the region of the initial dsRNA trigger. In contrast, when selecting for reads matching the targeted construct, we consistently found that they were derived exclusively from the region of the dsRNA construct (**Figure 2I**), and did not show any strand bias (**Figure 2J)**. Additionally, even though some SNPs were present in the long RNAseq reads matching the targeted genes, the small RNA sequences always matched the nucleotide sequence of the dsRNA construct (**Suppl Figure S4I**), suggesting that the cellular transcriptome is not used as a significant template for small RNA generation.

These findings thus indicate that dsRNA is effectively distributed and processed in *S. mediterranea*, but no widespread amplification of the small RNAs is detected. While some RNAi-induced phenotypes can be observed over long timescales, the direct effect of dsRNA on its mRNA target typically lasts only a few weeks. The observed dynamics are plausible without invoking a mechanism of siRNA or dsRNA amplification, and no evidence for such an amplification mechanism was uncovered.

### Systemic RNAi requires cycling stem cells

Over the course of our experiments exploring the requirements for planarian RNAi we noticed a striking phenomenon: lethally irradiated animals, that no longer contained stem cells, were ineffective at mounting a systemic RNAi response. Injection of dsRNA targeting the muscle gene *collagen-2*, into unirradiated animals resulted in a strong decrease in *collagen-2* mRNA as determined by qPCR (**Figure 3A**). However when the same dsRNA was injected into animals that had been depleted of neoblasts (**Suppl Figure 5A,B)**, no reduction in *collagen-2* mRNA was detected. A similar effect was detected for the knockdown of the eye gene *opsin*, which was highly effective in unirradiated animals, but was ineffective after lethal irradiation (**Figure 3B**). These findings suggested that the presence of stem cells might be required for systemic RNAi.

**Figure 3.**
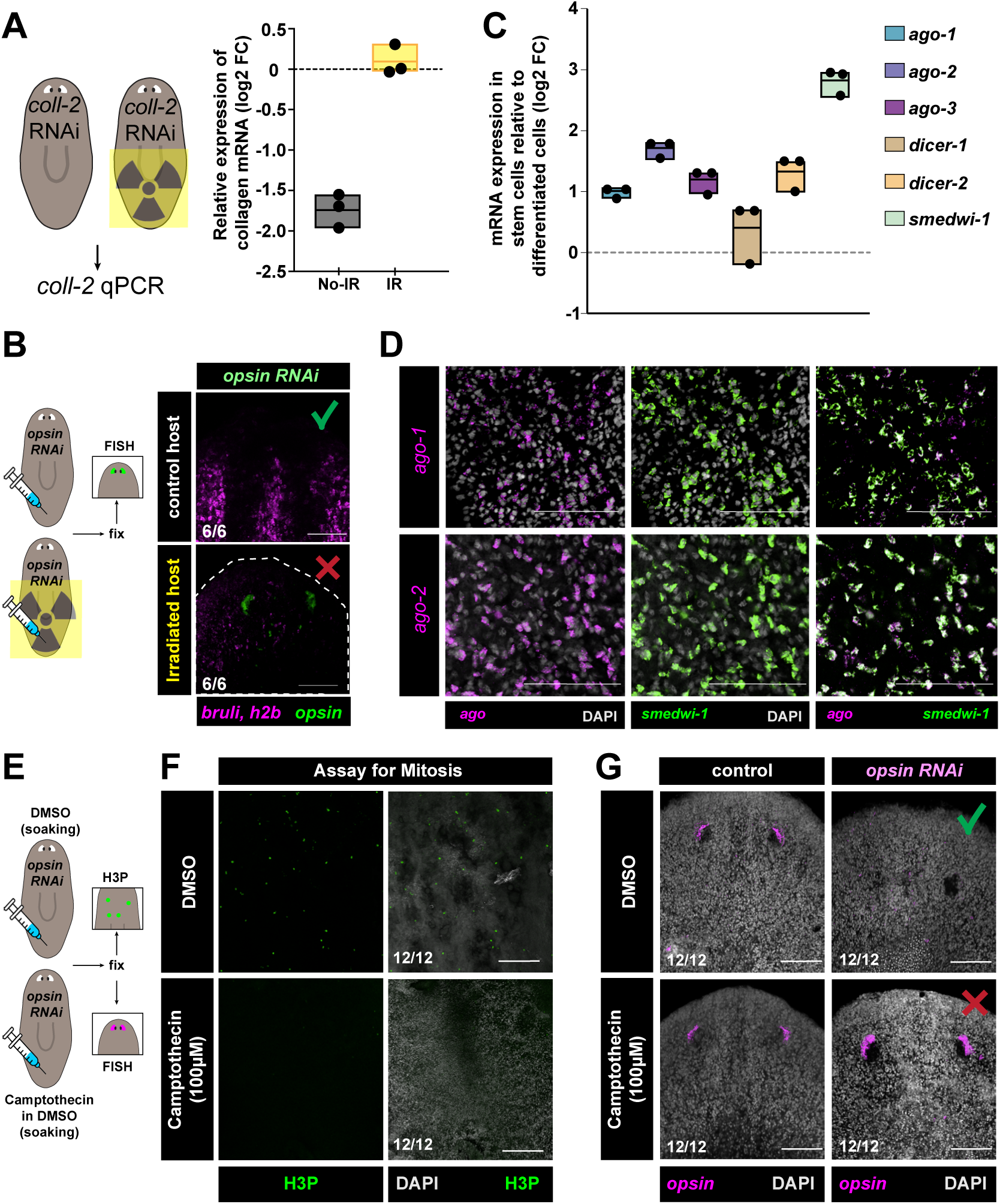
Systemic RNAi requires cycling stem cells **A.** Experimental schematic (left), and box plot (right) showing the change in *collagen-2* RNA level (log2 fold change relative to t=0) upon injection of *collagen-2* dsRNA in untreated or neoblast-depleted (irradiated, IR) animals. **B.** Confocal FISH images showing expression level of *opsin* (green) and neoblast genes *bruli* and *histoneH2B* (magenta) upon injection of *opsin* dsRNA in untreated or neoblast-depleted (irradiated, IR) animals. Green check marks indicate effective RNAi; red crosses indicate ineffective RNAi. **C.** Box plot showing the expression levels of Argonaute and Dicer genes and neoblast marker *smedwi-*1 in neoblasts compared to whole worms (log2 fold changes) as determined by qPCR. **D.** FISH images showing expression of Argonaute *ago-1* or *ago-2* (magenta) relative to neoblast genes *smedwi-1* (green). **E.** Experimental schematic for camptothecin treatment. **F.** Immunofluorescence of the mitotic marker H3P in control treated and camptothecin treated animals. **G.** FISH for *opsin* in control treated and camptothecin treated animals upon control dsRNA injection or injection of *opsin* dsRNA. In the presence of camptothecin, cell cycle is arrested, and RNAi-mediated knockdown is ineffective. Green check marks indicate effective RNAi; red crosses indicate ineffective RNAi.

We wondered whether some of the RNAi factors might be specifically expressed in the neoblasts. Transcripts for all three Argonautes were low in abundance in RNAseq datasets, but qPCR analysis of isolated neoblasts and differentiated cells revealed that all three were enriched in the neoblasts over the differentiated tissues (**Figure 3C**). FISH analysis confirmed localization of *ago-2* mRNA to stem cells (in agreement with (Rouhana et al. 2010; Li et al. 2011)), whereas *ago-1* mRNA, though still enriched in the neoblasts, was more broadly expressed (**Figure 3D**).

Transcript levels of *ago-3* were too low to obtain a clear signal by FISH. qPCR analysis also showed that expression of the miRNA-specific *dcr-1* was ubiquitous, but the transcript for the siRNA-specific *dcr-2* was similarly enriched in the neoblasts (**Figure 3C**). Together, these data show that the factors involved in the generation of siRNAs are indeed enriched in the neoblasts, which may explain the requirement for these cells.

Finally, since neoblasts are continuously cycling, we asked whether systemic RNAi depends simply on the presence of stem cells, or specifically on their proliferating activity. To test this, we treated animals with the cell cycle inhibitor camptothecin, which arrests neoblast division without eliminating the stem cell population (**Figure 3E-G, Suppl Figure S5C,D**). Remarkably, this treatment almost completely abolished RNAi efficiency, demonstrating that the cycling activity of stem cells, rather than their mere presence, is required for the systemic spread of the RNAi signal.

### Systemic RNAi is mediated by the spread of RNPs

The existence of systemic RNAi in planarians is well established, and our data (see above) indicate that the neoblasts play an important role in this process, but how the spreading of the silencing between cells works has remained unclear. In *C. elegans*, which has powerful systemic RNAi, the spreading agent is the dsRNA, and the dsRNA-binding membrane protein SID-1 is required cell autonomously for dsRNA uptake into the somatic cells (Winston et al. 2002). *S. mediterranea* encodes three homologs of SID-1, that based on publicly availably single cell sequencing data are enriched in cathepsin cells and glia (Fincher et al. 2018) (**Suppl Figure**

**S5E**). AlphaFold-based predictions of these SID-1 homologs indicate that planarian proteins form structures resembling those of *C. elegans* SID-1, suggesting a similar role in dsRNA import (**Suppl Figure S5F**). Indeed, we found that import of parenchymally injected long dsRNA into specific cells depended on these SID-1 homologs (**Suppl Figure S5G,H**). In contrast, dsRNA that was cut into ∼22nt fragments by RNaseIII did not translocate into cells upon injection, and did not induce silencing of a target gene, consistent with the expected specificity of SID-1 for long dsRNA (**Suppl Figure S5I**). Further, knockdown of *sid-1* resulted in the loss of dsRNA-mediated silencing of other genes, such as the eye gene *opsin*, and the neoblast genes *smedwi-1*, *smedwi-2*, and *histone-3* (**Figure 4A,B, Suppl Figure S5J**), indicating that SID-1 is essential for RNAi in *S. mediterranea*.

**Figure 4.**
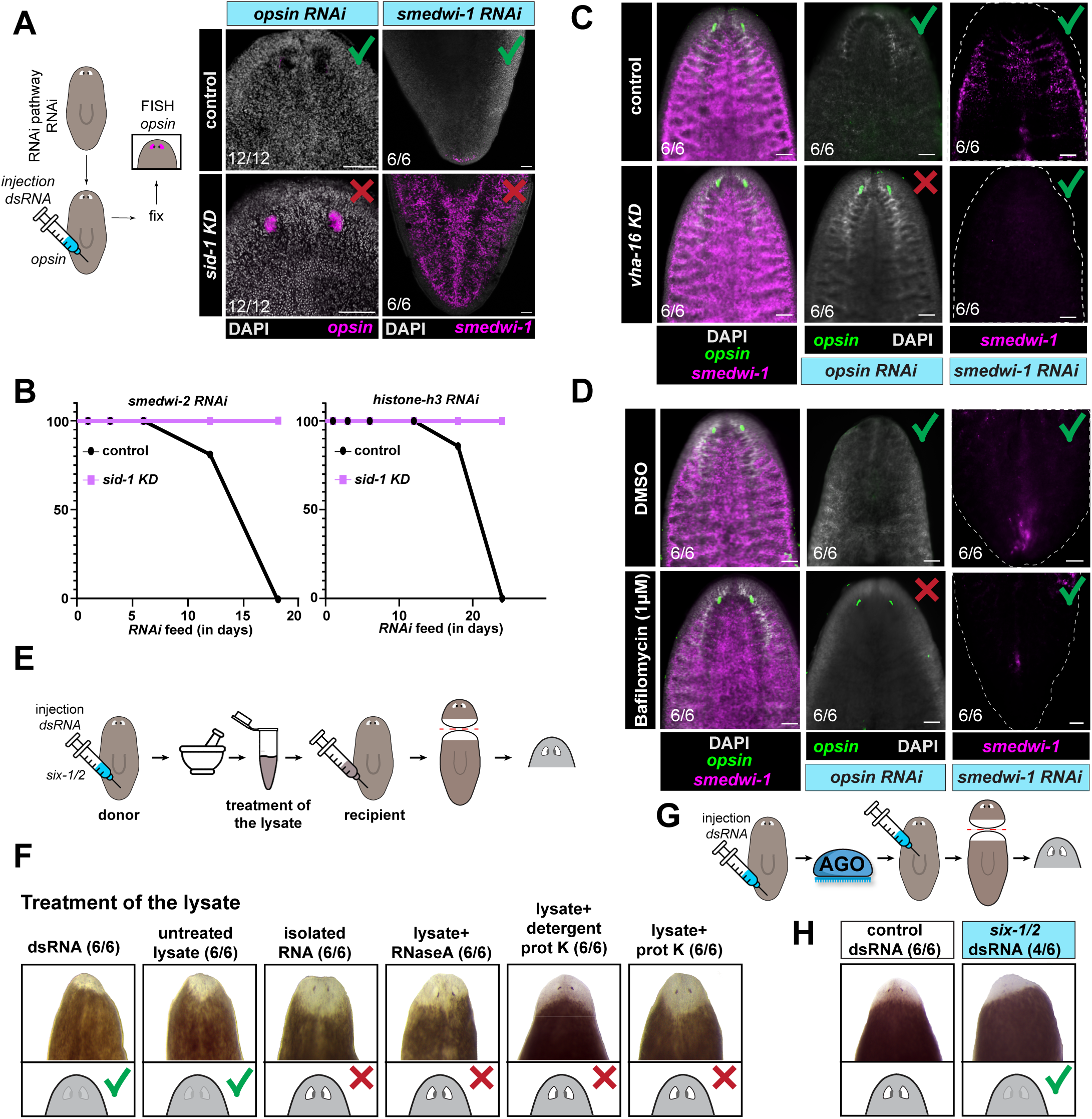
Systemic RNAi is mediated by the spread of RNPs **A.** Experimental schematic (left), and confocal FISH images (right) showing the efficiency of RNAi-mediated knockdown of eye transcript *opsin* (left) or neoblast transcript *smedwi-1* (right) in control animals (top), or upon knockdown of *sid-1* (bottom). Absence of SID-1 abolishes RNAi in both eyes and neoblasts. **B.** Survival curves of animals treated with dsRNA for neoblast genes *smedwi-2* or *histoneH3* show lethality in control animals, but continued survival in the absence of SID-1, indicative of ineffective RNAi. **C.** Confocal FISH images showing the efficiency of RNAi-mediated knockdown of eye transcript *opsin* (green) or neoblast transcript *smedwi-1* (magenta) in control animals (top), or upon knockdown of *vha-16* (bottom). Absence of VHA-16 abolishes RNAi in eyes but not in neoblasts. **D.** Confocal FISH images showing the efficiency of RNAi-mediated knockdown of eye transcript *opsin* (green) or neoblast transcript *smedwi-1* (magenta) in control animals (top), or upon treatment with Bafilomycin (bottom). Bafilomycin abolishes RNAi in eyes but not in neoblasts. **E.** Experimental schematic for the RNAi transfer experiments shown in F. **F.** Brightfield images and schematics of the images showing macroscopic eyes in newly regenerated heads upon injection of untreated or treated lysate from animals treated with dsRNA for eye gene *six-1/2*. Whereas untreated lysate transmits the silencing, treatment of the lysate with protease eliminates the silencing. **G.** Experimental schematic for the AGO transfer experiment shown in H. **H.** Brightfield images and schematics of the images showing macroscopic eyes in newly regenerated heads upon injection of isolated AGO complexes from animals treated with dsRNA for eye gene *six-1/2*. Green check marks indicate effective RNAi; red crosses indicate ineffective RNAi.

SID-1 was initially thought to form a channel through the cell membrane that would allow for the specific import of dsRNA (Shih and Hunter 2011). Recent structural studies of SID-1 however demonstrated that this protein rather is a dsRNA binding receptor that can enhance endocytosis of dsRNA into the cell (Zhang et al. 2024). Indeed, we found that in addition to SID-1, the gene *vha-16* which is involved in endocytosis in *C. elegans* and *Drosophila* (Saleh et al. 2006) was also required for systemic RNAi in planarians (**Figure 4C**). Furthermore, the endocytosis-inhibiting drug Bafilomycin also inhibited systemic RNAi (**Figure 4D, Suppl Figure S5K**). Remarkably, dsRNA-mediated silencing of stem cell genes was not sensitive to absence of *vha-16* or to bafilomycin, suggesting that the planarian neoblasts can use a different mode of dsRNA uptake, even though SID-1 was still required (**Figure 4A-D**).

RNAi initiated by dsRNA that enters through the intestine or through the body cavity, spreads through the entire planarian body and can affect gene expression in every cell in the body, even affecting new cells that were not present at the time of the original dsRNA trigger (**Figure 2C,G**). To gain further understanding of the molecular mediators of systemic RNAi, we took advantage of the sensitive nature of the eye regeneration (**Figure 2D**). We induced RNAi by injecting dsRNA against *six-1/2* or a control sequence in donor animals. On day 3 after injection, we made lysates from the injected animals and isolated several components for transplantation to recipient animals. Recipients were subsequently amputated to induce head regeneration and detect whether the transplanted material affected the regenerating eyes (**Figure 4E**). Crude lysates from *six-1/2* dsRNA-treated donors (RNA-∼1µg and Protein-∼100µg), but not control-treated donors, robustly induced RNAi in hosts (**Figure 4F, Suppl Figure S5L,M**). Isolated RNA from lysate was ineffective at transferring the RNAi effect, indicating that the transferring agent is not just the dsRNA or the siRNA. Remarkably, lysates treated with RNaseA or Proteinase K (either in the presence or in the absence of detergent) also did not transfer the RNAi effect (**Figure 4F, Suppl Figure S5M**), suggesting that both a protein and an RNA component are involved. Further, this experiment indicates that the transferable agent is not membrane-protected in our lysates. Based on these findings, we wondered whether the transferable agent might be an siRNA-loaded AGO protein. To test this, we isolated AGO complexes from the donors by the TraPR anion-exchange procedure (Lau et al. 2006) and transplanted these to hosts (**Figure 4G**). This effectively transferred the RNAi effect, indicating that this RNP is the most likely candidate for the transferable agent of systemic RNAi.

### Systemic RNAi requires siRNA biogenesis for initiation, and AGO-3 in the recipient cells

We next aimed to identify which of the biogenesis factors were involved in the initiation of the RNAi, and which were required for the spreading. To test whether the stem cells were required for the production of the transferable RNAi signal, we employed the same donor-host paradigm, but prior to the initial dsRNA injection, some donor animals were lethally irradiated, while others were left unirradiated (**Figure 5A**). Lysates from unirradiated donors efficiently transferred RNAi to recipient animals, whereas lysates from irradiated donors were ineffective, indicating that the stem cells are required for the initiation of the RNAi (**Figure 5B**).

**Figure 5.**
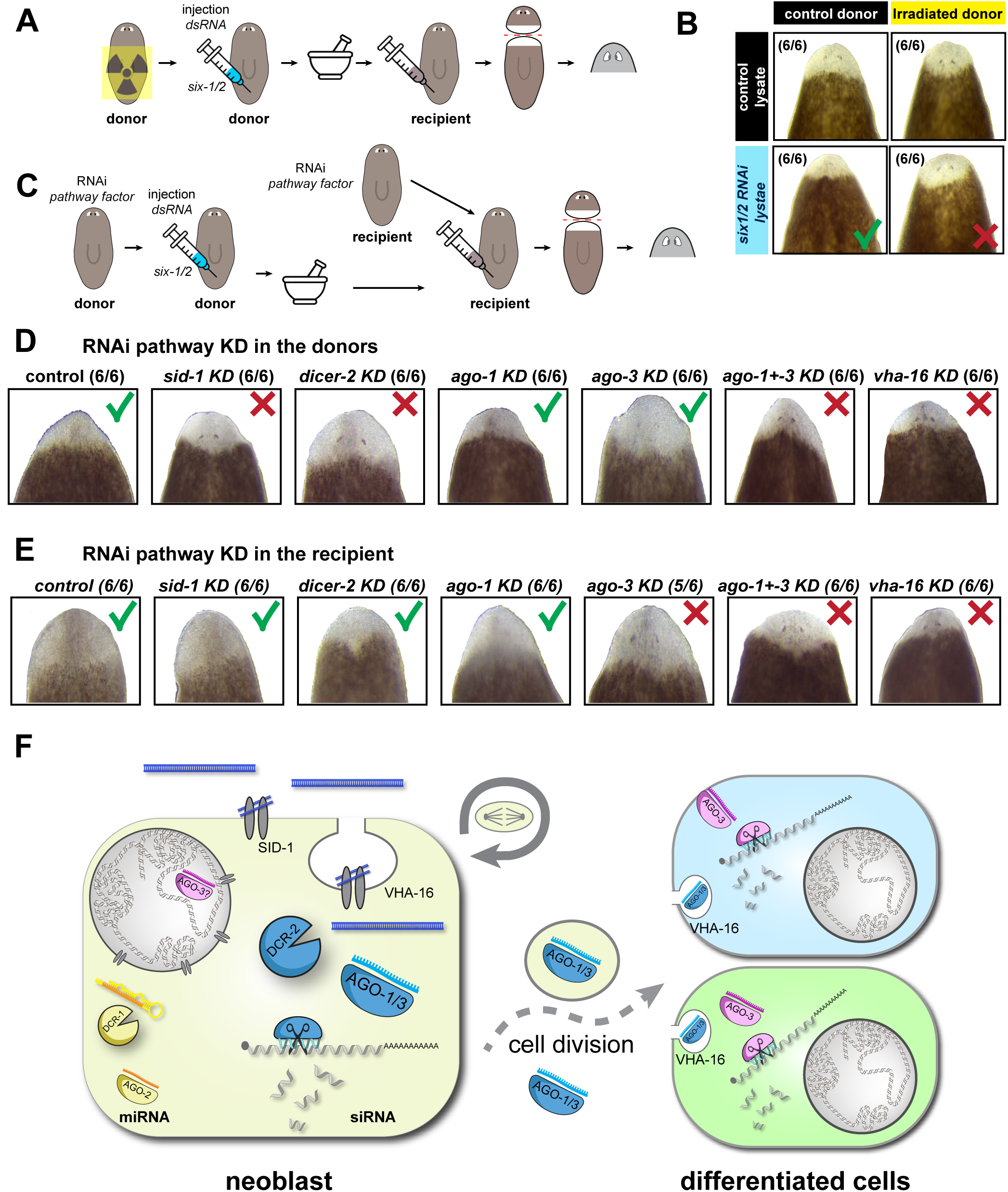
Systemic RNAi requires siRNA biogenesis in stem cells **A.** Experimental schematic for the irradiation and RNAi transfer experiment shown in B. **B.** Brightfield images showing macroscopic eyes in newly regenerated heads of recipient animals upon injection of lysate from unirradiated or irradiated animals treated with dsRNA for eye gene *six-1/2* or control dsRNA. **C.** Experimental schematic for the RNAi transfer experiment shown in D and E. **D.** Brightfield images showing macroscopic eyes in newly regenerated heads of recipient knockdowns in RNAi pathway factors upon injection of lysate from animals treated with dsRNA for eye gene *six-1/2*. **E.** Brightfield images showing macroscopic eyes in newly regenerated heads of recipients upon injection of lysate from donor knockdowns in RNAi pathway factors treated with dsRNA for eye gene *six-1/2*. Green check marks indicate effective RNAi; red crosses indicate ineffective RNAi. **F.** Model of planarian systemic RNAi. RNAi can be initiated from dsRNA injected into the parenchymal space. Silencing of transcripts in differentiated cells involves a two-step process that requires the neoblasts (planarian stem cells). In the first phase, the dsRNA binding membrane protein SID-1 is required, as well as the vesicle trafficking gene *vha-1*. Further, the processing requires an siRNA-specific Dicer, DCR-2, and the Argonaute proteins AGO-1 and AGO-3. Established RNAi can be transferred to a recipient animal through AGO-RNA complexes that may travel inside or outside of vesicles between cells. Uptake in differentiated cells requires the vesicle trafficking gene *vha-1*, and silencing requires the expression of the Argonaute AGO-3, suggesting that small RNA derived from the imported complex is handed off to cell-endogenous AGO-3 protein for targeting activity.

We next tested the requirement for the various proteins in the donors by treating donors with RNAi for each of these factors for 1 week before initiating RNAi against *six-1/2* by dsRNA injection (**Figure 5C**). We found that all factors that were required for the efficient induction of RNAi in the donor (SID-1, VHA-16, AGO-1+3 and DCR-2) were also required for the generation of the transferable agent (**Figure 5D**). When we however inverted the experiment and tested for the requirement of these factors in the host, we found that dsRNA interacting proteins SID-1 and Dcr-2 were no longer required for propagation of the RNAi, confirming the notion that dsRNA is not the transferred agent (**Figure 5E**). Rather than the redundant requirement for AGO-1 and AGO-3, during propagation AGO-3 was the only required Argonaute protein.

Further, we found that Bafilomycin still blocked transfer and that VHA-16 remained required (**Figure 5E**), indicating that some form of endo-or exocytosis likely is involved.

Together, our data show distinct requirements for the initiation phase and the transfer of RNAi (**Figure 5F**). Initiation requires the presence of cycling neoblasts, and the dsRNA binding protein SID-1 as well as some form of vesicle processing. It also requires the dsRNA slicing enzyme DCR-2 and Argonautes AGO-1 and/or AGO-3. This phase probably produces a mobile factor, that based on our experiments most likely is an AGO-siRNA complex. Transfer to other cells no longer requires factors that interact with dsRNA but still requires AGO-3 and a form of vesicle processing.

## Discussion

RNAi has an important and conserved role in the defense against viruses and mobile genetic elements. While many organisms have dedicated cells to recognize and eliminate infecting agents, activation and migration of such cells to the site of infection takes time. RNAi is rapid and cell autonomous, and thus can form a first line of defense for an infected cell, halting invading elements as soon as they initiate replication. As infecting agents are typically mobile and can spread to neighboring cells, it would be beneficial for the defending RNAi to spread to surrounding cells as well. Indeed, spreading of RNAi-mediated silencing between cells, known as systemic RNAi, has been described in multiple organisms (Jose and Hunter 2007). In plants, siRNAs were found to travel from cell to cell through plasmodesmata, as well as long-distance through the phloem (Molnar et al. 2010; Melnyk et al. 2011). In *C. elegans* systemic RNAi has been extensively studied, and involves mobile dsRNA that is internalized with the help of the dsRNA binding transmembrane protein SID-1 (Winston et al. 2002; Jose et al. 2011). In *Drosophila*, no mobile dsRNA or SID-1 is found, but RNA silencing components have been proposed to spread from cell to cell through nanotubes (Karlikow et al. 2016). And while in mammalian systems dsRNA mediated silencing is thought to be less impactful as it would activate interferon signaling, extracellular small RNAs have been detected, and may have signaling functions (Hoy and Buck 2012; Fritz et al. 2016).

Here, we find that systemic RNAi in planarians likely involves the import of dsRNA from the extracellular space into the cells, as it requires SID-1, similar to the situation in *C. elegans*. SID-1 was expressed in many of the planarian cell types, suggesting that extracellular dsRNA could reach the cytoplasm of most cells. Surprisingly, we found that the absence of active neoblasts however led to ineffective silencing. This finding has two implications: first, it suggests that there must be something specific about the neoblasts that allows them (and not other cells) to initiate silencing; and second, it suggests that there must be a mobile RNAi intermediate that migrates from the neoblasts to the differentiated cells.

Our experiments revealed that the mobile RNAi intermediate that migrates from the neoblasts to the differentiated cells, most likely is an Argonaute-siRNA complex. To address the nature of the mobile RNAi intermediate, we initiated RNAi in donor animals and transferred various components to a recipient animal to evaluate their ability to transfer silencing. We found that the transferrable silencing agent was sensitive to both RNase and protease, and that the transfer of isolated RNA was insufficient, arguing against dsRNA or siRNA as a transferring agent. We further found that SID-1 and DCR-2 were not required in the recipient animals, confirming the notion that the second mobile agent is distinct from the dsRNA trigger. Finally, purified AGO complex from the donor animals was able to transfer silencing. These data all converge on the notion that AGO-RNA complexes can be imported into differentiated cells to induce silencing.

Our RNAi transfer experiments revealed that the silencing in recipient cells was independent of siRNA biogenesis factors DCR-2 and AGO-1, but still required expression of AGO-3. Our sequencing data, qPCR data, and FISH, had indicated that *ago-3* is very low expressed, making it difficult to imagine how this protein could accomplish a substantial silencing effect in every planarian cell. Its expression however was significantly increased upon exposure to dsRNA, indicating that this protein may be generated on demand. Additionally, the N-terminal region of AGO-3 contains a predicted Nuclear Localization Signal, suggesting that loaded AGO-3 may be imported into the nucleus where a small amount of protein could have a meaningful effect on the transcriptional outcome. The requirement for AGO-3 in the recipient animals further suggests that the mobile AGO-RNA complex does not directly effectuate silencing in the recipient cells but rather loads cell-autonomous AGO-3. We did not find any evidence for amplification of the RNA involved in RNAi-mediated silencing, meaning that the RNA loaded onto AGO-3 must have been part of the imported AGO-RNA complex. Interestingly, a similar hand-off mechanism was recently proposed in HeLa cells (Ghoshal et al. 2021). AGO-1 was required in the donor cells but not in the recipients, indicating that it is likely not effective at silencing and thus may rather be involved in the transport of the RNAi intermediate. AGO-1 is characterized by a somewhat divergent N-terminal region and PAZ domain. The N-terminal domain of AGO proteins is thought to be important for the unwinding of the RNA duplex after loading (Kwak and Tomari 2012). It is possible that the N-terminal domain of AGO-1 is less effective in this process, and that AGO-1 therefore mostly binds dsRNA, increasing the amount of RNA that is imported into the recipient cells. Additionally, the poorly conserved PAZ domain may result in less stable binding of the RNA, facilitating a potential hand-off to AGO-3. Further details of this mechanism will have to be investigated in future studies.

Systemic RNAi often involves some form of vesicle-mediated transport. A screen for genes required for the uptake of dsRNA in *Drosophila* S2 cells primarily identified factors involved in endocytosis (Saleh et al. 2006; Ulvila et al. 2006), and similar factors were found to affect the efficiency of systemic RNAi in *C. elegans* (Tijsterman et al. 2004; Saleh et al. 2006). Initially, this involvement of a vesicle-based mechanism was surprising. However, the recent finding that SID-1 represents a dsRNA receptor rather than a channel, proposes a new model for dsRNA uptake that involves endocytosis of SID-1-bound dsRNA, thereby explaining the involvement of these factors (Hinas et al. 2012; Jose et al. 2012; McEwan et al. 2012; Zhang et al. 2024). In planarians, we found that the vacuolar ATPase *vha-16*, which is involved in endocytosis and related vesicle trafficking (Mellman et al. 1986; Futai et al. 2019), was required for silencing in differentiated cells, and correspondingly, that the vacuolar ATPase inhibitor bafilomycin-A1 inhibited systemic RNAi. Remarkably, *vha-16* and bafilomycin did not affect RNAi in the neoblasts, suggesting that exactly these cells that initiate the silencing response from exogenous dsRNA use a different mechanism of dsRNA import. The vesicle-related process in planarians thus appears to primarily concern the second mobile agent, the AGO-RNA complex. Given that *vha-16* was required both in the donor animals and in the recipient cells, it appears that this vesicle-based process is involved in both the dispersion and reception of the AGO-RNA complexes.

Our finding that injection of purified AGO-RNA complexes was sufficient to transfer silencing indicates that encapsulation by a membrane is not required to accomplish spread, import, and silencing in differentiated cells. Recent studies in Arabidopsis and human plasma revealed that transported siRNAs and miRNAs in plants and humans are largely localized outside of vesicles (Arroyo et al. 2011; Baldrich et al. 2019; Jeppesen et al. 2019; Geekiyanage et al. 2020; Tosar et al. 2021; Zand Karimi et al. 2022; Jeppesen et al. 2023), confirming the notion that such complexes do not require the protection of a membrane in the extracellular space. The most parsimonious model thus is that AGO-RNA complexes synthesized in the neoblasts are excreted by exocytosis and taken up by endocytosis in the recipient differentiated cells, but the involvement of lipid components cannot be excluded.

A puzzling aspect of planarian RNAi has been its efficiency in the absence of recognizable RNA dependent RNA polymerases (RdRPs). The fact that feeding or injection of dsRNA can produce sufficient amounts of silencing complexes to affect every cell of the planarian body has led to the notion that some other amplification mechanism must be in place. While it is not possible to definitively exclude the existence of such a mechanism in planarians, we found no evidence to support its involvement in typical experimentally induced RNAi. We found no evidence of siRNAs from regions other than the introduced dsRNA construct, and we only recovered siRNAs with the exact sequence of the exogenously introduced dsRNA, even when that was distinct from the genomic sequence. Further, by comparing the number of dsRNA molecules introduced (∼10^10^ molecules per animal) to the number of cells in the planarian body (∼10^5^ cells per animal), it appears that no amplification needs to be invoked to allow for systemic distribution.

Knowledge of the processes and mechanisms involved in RNAi is important for the accurate interpretation of the outcomes of planarian knockdown experiments, but moreover, it is instrumental for our understanding of the defense mechanisms that organisms employ to protect themselves from mobile genetic elements. Such defenses are crucial for the maintenance of genome integrity and thus have major impact on organismal survival and evolutionary processes. Furthermore, understanding how RNA-based silencing is processed and spreads between cells can provide important insights for the development of agricultural and therapeutic applications. Several commercial RNAi-based strategies have been developed to reduce the damage to crops by herbivorous arthropods or pathogens (Wytinck et al. 2020; Duanmu et al. 2025; Zhong et al. 2025). And several drugs targeting liver cells by RNAi-based silencing are already approved for clinical application (Friedrich and Aigner 2022; Corydon et al. 2023). Major challenges however remain concerning the uptake and systemic spread of the silencing agents, and improved insights in these processes thus could inspire the formation of more effective formulations. Our finding provide several new ideas. We find that activation of specific cell types may enable vastly enhanced silencing, while AGO proteins still need to be present in the targeted cells. Further our data indicate that AGO-RNA complexes could be conserved mobile agents in the communication between cells, while still requiring a vesicle-based process for import. These insights will inform further studies to identify analogous processes, and eventually may lead to new formulations and improved application strategies.

## Methods

### Planarian culture and animal husbandry

Asexual *Schmidtea mediterranea* clonal strain CIW4 and sexual strain S2 were maintained as previously described (Newmark and Sanchez Alvarado 2000). Animals were cultured in 1× Montjuïc salts at 20°C, fed homogenized beef liver paste every 1-2 weeks, and expanded through repeated cycles of amputation or fission and regeneration. Animals were starved for 1 week prior to all experiments. Unless stated otherwise, experiments were performed using asexual animals. For experiments requiring sexual reproduction, *Schmidtea polychroa* was maintained under standard laboratory conditions and used for crossing assays as described below.

### Sublethal and lethal irradiation

Animals were exposed to gamma irradiation using a MultiRad350 irradiator (Precision X-ray) equipped with a copper filter. For partial stem cell impairment, animals were irradiated with 500 Rads (low dose) or 1500 Rads (medium dose) and used for experiments 3 days post-irradiation. For complete stem cell ablation, animals received a lethal dose of 3000 Rads and were analyzed 6 days post-irradiation, a time point at which neoblasts are fully depleted.

Following irradiation, animals were maintained in planarian water supplemented with gentamicin (50 μg/ml).

### Pharmacological treatments

To inhibit neoblast proliferation without eliminating the stem cell population, animals were soaked in camptothecin (100 μM) in the presence of 0.1% DMSO to induce cell cycle arrest. Effective inhibition of neoblast proliferation was confirmed by loss of phospho-Histone H3-positive cells. Animals were washed extensively in fresh planarian water prior to dsRNA injection or subsequent RNAi assays.

For inhibition of vesicle trafficking and endocytosis, animals were treated with Bafilomycin A1 (1µM) prior to dsRNA exposure or RNAi analysis.

### dsRNA synthesis and RNA interference

Gene fragments of 0.5-2 kb were amplified from planarian cDNA using gene-specific primers containing adaptor sequences (Supplementary Table S1). PCR products were cloned into pGEM-T (Promega) and verified by Sanger sequencing. Sense and antisense RNAs were synthesized in vitro using T7 RNA polymerase from PCR templates containing flanking T7 promoter sequences (TAATACGACTCACTATAGG), annealed at 37°C for 30 min, and verified by gel electrophoresis.

For RNAi by feeding, (∼1µg) dsRNA was mixed with homogenized beef liver paste mixed with food coloring and fed to animals at 3-day intervals. Feeding regimens varied by target gene as indicated in figure legends. dsRNA corresponding to *C. elegans unc-22* was used as a negative control.

For RNAi by injection, dsRNA was diluted 1:10 in water and injected into the planarian parenchyma or body cavity using pulled glass capillaries of 0.02 inches inner diameter (Drummond™ Capillaries for Nanojet II™ Injectors, Fisher Scientific). The amount of injected dsRNA was ∼0.1 *p moles* unless indicated otherwise, and animals were analyzed at defined time points following injection. Regeneration experiments were initiated by either head, trunk or tail amputations at specified days post-injection or post *RNAi* as indicated.

### Whole-mount fluorescent in situ hybridization and immunofluorescence

Whole-mount fluorescent in situ hybridization (FISH) and immunofluorescence were performed as previously described (King and Newmark 2013). Briefly, animals were fixed in formaldehyde, bleached, treated with proteinase K (2 μg/ml), and hybridized overnight at 56°C with DIG-, or fluorescein-labeled riboprobes. Signal detection was performed using HRP-conjugated antibodies and tyramide signal amplification. Specimens were counterstained with DAPI. Immunofluorescence staining used rabbit anti-phospho-Histone H3 (Ser10; 1:750), mouse, followed by HRP-conjugated secondary antibodies and tyramide development.

### Quantitative PCR (qPCR) analysis

Total RNA was isolated using TRIzol (Invitrogen) and quantified using a Qubit fluorometer. cDNA was synthesized using oligo(dT) primers by ProtoScript II Reverse Transcriptase (NEB) from 1 μg RNA. Gene-specific RT primers used are included in Supplementary Table S1. qPCR reactions were performed using EvaGreen Master Mix (Biotium) on a QuantStudio 3 Real-Time PCR System (Applied Biosystems). Relative expression levels were calculated using ΔΔCt normalization to internal reference genes. For each experiment, all reactions were performed in parallel on the same plate.

### Quantification of small RNAs by poly(A)-tailing RT-qPCR

Small RNA abundance was measured using a polyadenylation-based reverse transcription quantitative PCR (qPCR) approach. This method involves enzymatic polyadenylation of small RNAs, reverse transcription using an anchored oligo-dT primer containing a 5′ adapter sequence, and qPCR amplification using an adapter-specific reverse primer and a small RNA-specific forward primer. Small RNA-specific primers were tested for specificity and quantitative performance prior to use.

Total RNA was isolated as described above, and 1 µg RNA was polyadenylated using Poly(A) polymerase according to the manufacturer’s instructions. Polyadenylation reactions were incubated at 37°C for 30 min and then cooled on ice. Reverse transcription was performed using ProtoScript II reverse transcriptase (NEB) and an anchored oligo-dT-(V)_N_ primer containing a 5′ adapter sequence. Following reverse transcription, RNA templates were removed by RNase H treatment.

Quantitative PCR was performed using EvaGreen chemistry with a small RNA-specific forward primer and an adapter-specific reverse primer. Reactions were run using standard qPCR cycling conditions, and data were analyzed as described as above.

### Small RNA sequencing analysis

Raw small RNA sequencing reads were processed using fastp (version 0.21.0) for quality filtering and adapter trimming. Reads shorter than 18 nucleotides were discarded (--length_required 18) to remove degradation products and non-informative sequences, while an average Phred quality score threshold of 20 (--average_qual 20) was applied to ensure high-confidence base calls (Chen et al. 2018). These parameters are commonly used in small RNA sequencing studies to balance read retention and data quality.

Following preprocessing, reads were categorized based on length to distinguish small RNA classes. Reads shorter than 28 nucleotides were classified as siRNAs or miRNAs, whereas reads of 28 nucleotides or longer were classified as piRNAs. This threshold reflects the established size distributions of small RNAs in metazoans and has been widely adopted in previous small RNA profiling studies.

Filtered reads were aligned to the Schmidtea mediterranea reference genome (schMed3; (Ivankovic et al. 2024)) using STAR (version 2.7.11a) (Dobin et al. 2013). Alignments were performed allowing up to two mismatches per read to account for sequencing errors and biological variation while maintaining mapping specificity. A maximum of 100 alignments per read was permitted to retain reads mapping to repetitive genomic regions, which are characteristic of piRNA loci.

To prevent spliced alignments, which are not expected for small RNAs, genome indices were generated without annotation (GTF) files, and the STAR parameter --alignIntronMax 1 was applied. This approach ensures that only contiguous genomic alignments are reported and reduces spurious mappings.

### Neoblast isolation and analysis

Neoblasts in G2/M phase (X1), and differentiated cells (Xins) were isolated by fluorescence-activated cell sorting (FACS) based on Hoechst DNA content, as previously described. Purified cell populations were subsequently used for RNA extraction and quantitative PCR as indicated.

### Cell lysate preparation

Planarian cells were dissociated by finely mincing 12 medium-sized (5–7 mm) asexual Schmidtea mediterranea with a razor blade and incubating the tissue in 1× CMF buffer (150 mM HEPES, pH 7.3; 33 mM NaH PO; 95 mM NaHCO; 140 mM NaCl; 160 mM KCl; 0.24% glucose; filter-sterilized through a 0.2 µm vacuum filter). Tissue suspensions were rocked on a nutator for 5 min, followed by gentle trituration with a P1000 pipette for 10 min, repeated three times or until the tissue was visibly dissociated. Cells were pelleted by centrifugation at 300 × g (soft acceleration/deceleration) for 5 min, resuspended in 1 mL CMF buffer, and passed through a sterile 40 µm cell strainer (Corning).

Single-cell suspensions were lysed in cellular lysis buffer (20 mM Tris-HCl, pH 7.5; 150 mM NaCl; 1.5 mM MgCl; 0.6% Triton X-100; 0.5 mM DTT) supplemented with cOmplete™, Mini, EDTA-free protease inhibitor cocktail (Roche). For detergent-free lysates, Triton X-100 was omitted. Crude lysates were clarified by centrifugation at 1,000 × g at 4°C for 5 min. Protein concentrations were measured by Bradford assay using absorbance at 280 nm, and approximately 100 µg of total protein was used per injection. Where indicated, lysates were treated with RNase A (50 µg/mL; Sigma) or Proteinase K (20 µg/mL; NEB) at 37°C for 30 minutes prior to injection.

### Q-Sepharose-based enrichment of Argonaute (AGO) complexes

Planarian cell lysates were subjected to batch binding on Q-Sepharose anion-exchange resin to enrich AGO-associated small RNA complexes based on their differential salt-dependent binding properties. Animals were rinsed in Binding Buffer (20 mM HEPES-KOH, pH 7.9; 10% glycerol; 100 mM potassium acetate [KOAc]; 0.2 mM EDTA; 1.5 mM MgCl), freshly supplemented with 1 mM DTT and 1× EDTA-free protease inhibitor cocktail (Roche). Approximately 25 medium-sized planarians were snap-frozen in liquid nitrogen and mechanically disrupted using a mortar and pestle in 500 µL Binding Buffer. Lysates were further homogenized using a dounce and clarified by centrifugation at 12,000 × g for 10 min at 4°C. Supernatants were retained as clarified lysates.

Q Sepharose™ Fast Flow resin (Cytiva Life Sciences) was equilibrated at 4°C by washing twice with sterile nuclease-free water and three times with Binding Buffer. Clarified lysates were adjusted to a final volume of 1 mL in Binding Buffer and supplemented to 400 mM KOAc before binding to equilibrated Q-Sepharose slurry (∼0.75 mL packed resin per reaction) for 10 min at 4°C with gentle mixing. Following incubation, the resin which had bound ribosomes and most other RNPs was pelleted at 2,000 × g for 2 min, and the supernatant containing AGO proteins and associated small RNAs was collected for downstream applications.

### Donor-recipient RNAi transfer assays

To assess systemic RNAi transfer, donor animals were injected with dsRNA targeting six-1/2 or with control dsRNA. Three days post-injection, donor animals were processed to generate crude lysates as described above (∼1 µg RNA and/or ∼100 µg protein). Lysates were subjected to the indicated treatments: Proteinase K digestion or Proteinase K digestion in the presence of detergent (0.6% Triton X-100) at 37°C for 30 minutes, or purification of RNA by Trizol.

Untreated or treated lysates were transplanted into naïve (wild type) or RNAi knockdown recipient animals by injection, followed by head amputation to assay eye regeneration. RNAi transfer efficiency was evaluated by FISH and morphological analysis of regenerated eyes. For Argonaute–RNP transfer experiments, AGO-containing ribonucleoprotein complexes were isolated using Q Sepharose™ Fast Flow resin and injected into recipient animals.

### Testing sexual transfer of RNAi

*Schmidtea polychroa* worms were subjected to prolonged feeding (5 weeks) of *six-1/2* or control dsRNA to assess whether RNAi effects induced in the parents persist into the F1 generation through embryogenesis. Sixteen worms per condition were amputated and allowed to regenerate to verify the formation of macroscopic eyes. Starting three weeks post-amputation, they were fed∼1µg dsRNA twice weekly. Different food coloring was used for control or *six-1/2* animals to track treatment identity. Animals were maintained in gentamycin-supplemented planarian water starting one week prior to mating.

After 35 days of RNAi treatment, and after confirming the loss of macroscopic eyes in regenerating heads, the individual mating assays were initiated. *Six-1/2 KD* (n=8) worms were paired individually with control (n=8) worms, while the remaining worms were paired within the same treatment, yielding mixed (*six-1/2* × control) and same-treatment (*six1/2* × *six1/2* or control × control) crosses. Pairs were placed in fresh planarian water and allowed to mate undisturbed for two days. The worms were separated on day 3 and returned to individual dishes for tracking of capsules and for continued RNAi feeding. The capsules were collected in 12-well plates for individual tracking. Capsule and hatchling numbers were recorded to assess fertility and viability, and hatchlings were imaged to score eye phenotypes as a readout of *six-1/2* RNAi tranfer. Feeding, mating, and capsule collection were repeated for four consecutive weeks. To verify sustained RNAi in parental animals, heads were fixed at the end of the assay and processed to confirm target gene knockdown.

### Phylogenetic analyses and protein sequence alignments

Phylogenetic analyses of AGO and DICER proteins were conducted using selected full-length sequences listed in Supplementary Table 1. Full-length proteins were chosen to preserve conserved and divergent domains relevant for functional and evolutionary inference.

Protein sequences were aligned using MUSCLE (version 3.8.1551) with default parameters (Edgar 2004), which provide a reliable balance between alignment accuracy and computational efficiency for protein datasets of moderate size.

Phylogenetic trees were inferred using FastTree (version 2.1.11), which estimates approximately maximum-likelihood trees from multiple sequence alignments (Price et al. 2010). The parameters-spr 4-mlacc 2-slownni-gamma were applied to increase tree accuracy by optimizing subtree-prune-regraft moves, refining likelihood calculations, and modeling among-site rate heterogeneity. This parameter combination is well suited for resolving deep and shallow branching patterns in conserved protein families.

Tree visualization and annotation were performed using iTOL (Interactive Tree Of Life) (Letunic and Bork 2021).

### Protein Domain and Structure predictions

Protein sequences corresponding to S. mediterranea ago-1 (SMU15002348), ago-2 (SMU15038936), ago-3 (SMU15032988), dicer-1 (SMU15034558), and dicer-2 (SMU15033102) were predicted using orfipy (version 0.0.4) from the corresponding transcript sequences (Singh and Wurtele 2021).

Conserved protein domains were identified using InterProScan (version 5.61–93.0) with default parameters (Paysan-Lafosse et al. 2023), which integrate multiple domain databases to provide comprehensive functional annotation. Protein domain schematics were generated using drawProteins, enabling clear visualization of conserved and variable regions.

Protein structure predictions were performed using AlphaFold3 (Abramson et al. 2024). RNA molecules were included as ligands to reflect biologically relevant RNA–protein interactions. For AGO-1, AGO-2, and AGO-3, a 21-nucleotide RNA sequence (CCAGCUUUCAUAGUGAUCUCA) was used, corresponding to the canonical length of guide RNAs bound by Argonaute proteins.

DICER-1 was modeled using a pre-miRNA hairpin sequence (CCUUAUUAUAUUGCUGUCAAGUAAGCACUACUGGACGGAAGUGUAAUGAGGG) to reflect its role in processing stem–loop RNA substrates. DICER-2 was modeled using a 50-nucleotide double-stranded RNA substrate, provided as a sequence (GCCAACGAGUUCAGCCAAACGAUGGGCAUAAUGCGUUGGAACAGGAACUC) and its reverse complement as two separate RNA molecules. This length was chosen to approximate a short dsRNA intermediate representative of DICER-2 substrates.

All predicted structures were visualized and analyzed using PyMOL (Schrödinger, LLC).

### Microscopy and image quantification

Images were acquired using a Zeiss LSM800 confocal microscope under identical settings for control and experimental samples. Quantification of fluorescence intensity and cell counts was performed using Fiji (Schindelin et al. 2012) and PIQ software (Allikka Parambil et al. 2024).

## Statistical analysis

Statistical significance was assessed using two-tailed Student’s *t* tests in Prism (GraphPad). Genome-wide analyses were conducted using the associated bioinformatics tools as described above. All experiments were performed with at least three biological replicates unless otherwise stated.

## Resource availability

Sequencing data generated over the course of this study has been deposited in the SRA under accession number PRJNA1415216.

## Acknowledgements

We thank Max Bales and Colin Smith for experimental support, and thank all other members of the Van Wolfswinkel lab for support and discussion. We are grateful to the Keck DNA Sequencing Facility at Yale University for services provided. This work was supported by NIH grant R35GM158281 (to JCvW).

## Author contributions

Conceptualization, SAP and JCvW.; data acquisition, SAP, KAG, LC, and MW; methodology, AP and JCvW; formal analysis and software, AP and HL; funding acquisition, JCvW; supervision, JCvW.; writing – original draft, JCvW; writing – review and editing, SAP and JCvW.

