## Supplemental data for "Systemic RNAi in planarians depends on spread of RNPs from active stem cells"

### SUPPLEMENTAL FIGURES

**Supplemental Figure S1**

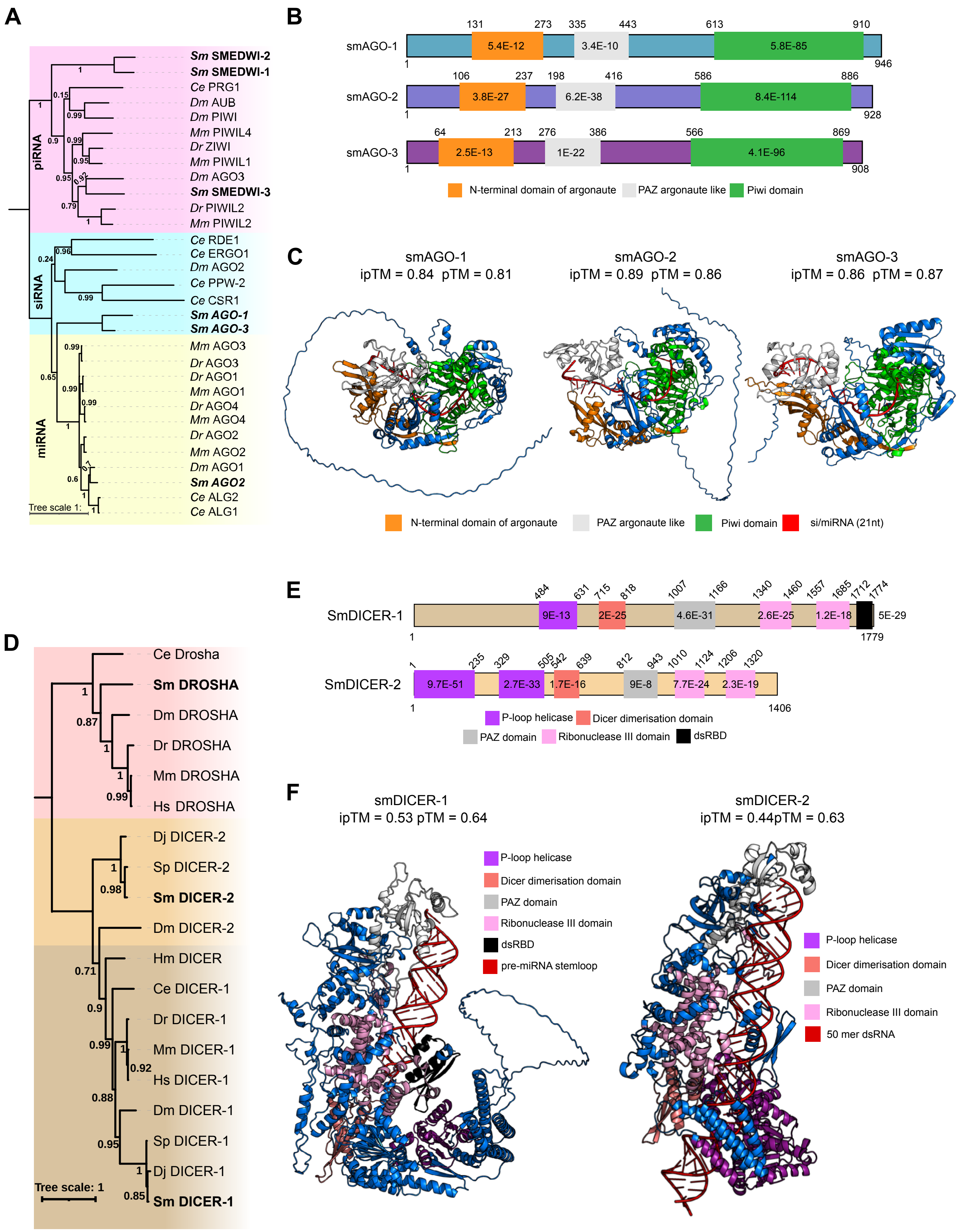

**Supplementary Figure S1. Planarian Argonaute and Dicer proteins.**

**A.** Phylogenetic tree of metazoan Argonaute (AGO) proteins, with *Schmidtea mediterranea* proteins indicated in bold. Branch numbers indicate bootstrap support. Species abbreviations: Ce, *Caenorhabditis elegans*; Dr, *Danio rerio*; Dm, *Drosophila melanogaster*; Hs, *Homo sapiens*; Mm, *Mus musculus*; Sm, *Schmidtea mediterranea*.

**B.** Predicted domain architecture of *S. mediterranea* AGO proteins based on InterProScan analysis (Paysan-Lafosse et al. 2023). Numbers within domains indicate InterProScan scores.

**C.** AlphaFold-predicted models of *S. mediterranea* AGO-1, AGO-2, and AGO-3 in complex with a 21-nt small RNA (red). pTM indicates confidence in the overall structure, and ipTM indicates confidence in the predicted protein–RNA interaction.

**D.** Phylogenetic tree of metazoan DICER proteins, with *S. mediterranea* proteins indicated in bold. Branch numbers indicate bootstrap support. Species abbreviations: Ce, *Caenorhabditis elegans*; Dr, *Danio rerio*; Dm, *Drosophila melanogaster*; Hs, *Homo sapiens*; Mm, *Mus musculus*; Hm, *Hofstenia miamia*; Dj, *Dugesia japonica*; Sp, *Schmidtea polychroa*; Sm, *Schmidtea mediterranea*.

**E.** Predicted domain architecture of planarian DICER proteins based on InterProScan analysis. Numbers within domains indicate InterProScan scores. The first helicase domain in SmDICER-1 and dsRBD in SmDICER-2 were not recognized by InterProScan but conserved residues were identified by alignment.

**F.** AlphaFold-predicted models of *S. mediterranea* DICER-1 and DICER-2 in complex with a pre-miRNA stem-loop or a 50-bp dsRNA substrate (red), respectively. pTM indicates confidence in the overall structure, and ipTM indicates confidence in the predicted protein–RNA interaction.

#### Supplemental Figure S2

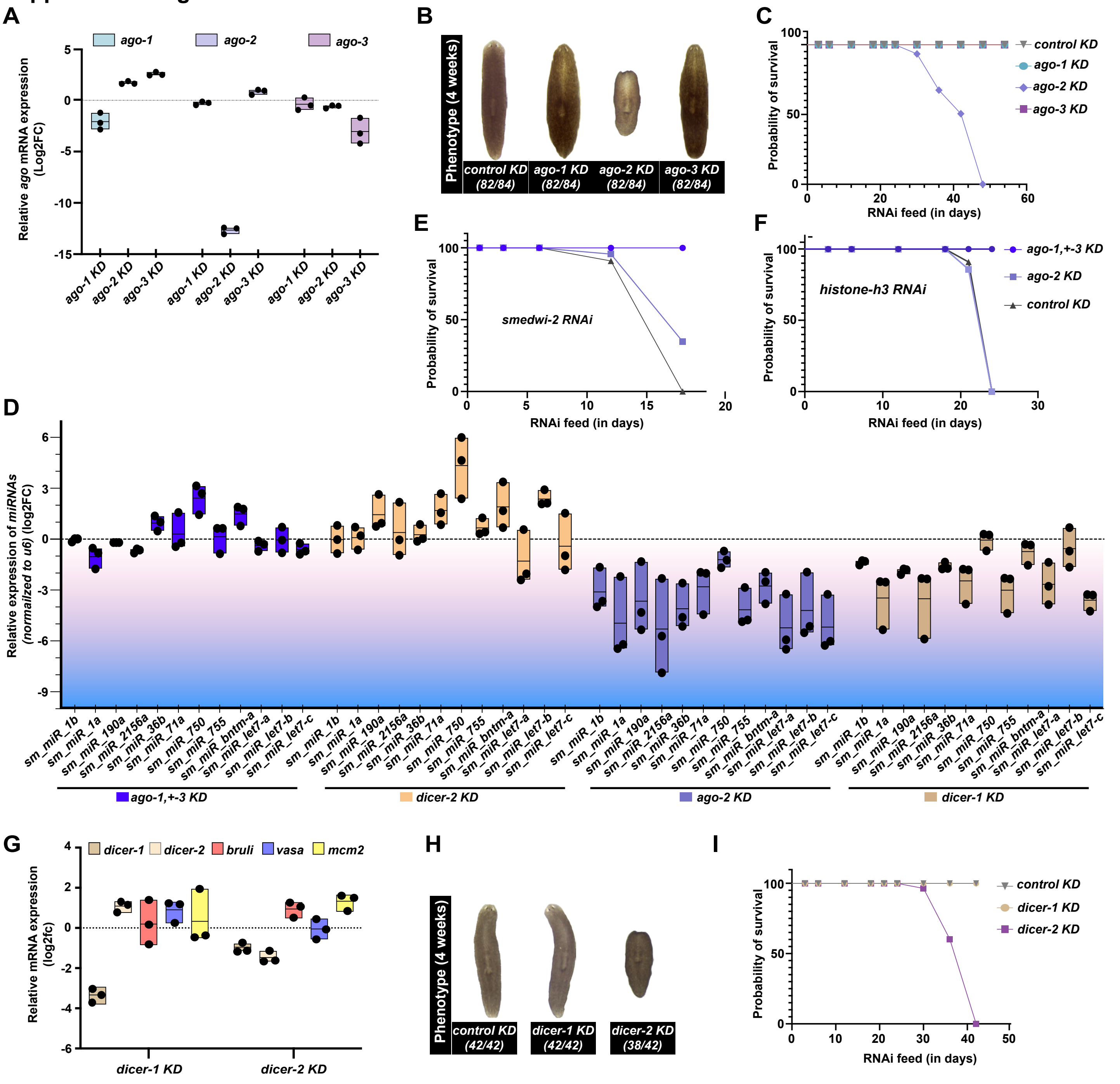

**Supplementary Figure S2. Distinct molecular requirements for siRNA and miRNA pathways in planarians.**
**A.** mRNA changes of argonaute family transcripts following the indicated knockdowns, as measured by qPCR.
**B.** Representative brightfield images of phenotypes following argonaute knockdown. *ago-2* RNAi causes progressive head regression, whereas *ago-1*, *ago-3*, and control animals remain morphologically normal.
**C.** Survival curve of animals treated with dsRNA targeting argonaute genes. *ago-2* knockdown is lethal, while *ago-1*, *ago-3*, and control treatments show sustained survival.
**D.** qPCR measurements corresponding to the miRNA heatmap in Fig. 2A, shown as log2 fold changes in miRNA abundance following knockdown of *ago-1+3*, *ago-2*, *dcr-1*, or *dcr-2* relative to controls.
**E.** Survival curve following RNAi treatment against the neoblast gene *smedwi-2* in control animals or argonaute knockdown animals. Control and *ago-2* animals exhibit the expected lethality upon exposure to *smedwi-2* dsRNA, while combined *ago-1+3* knockdown animals survive, consistent with impaired RNAi. Experiments were initiated after two weeks of argonaute knockdown to control for intrinsic *ago-2* lethality.
**F.** Survival curve following RNAi treatment against the neoblast gene *histoneH3* in control animals or argonaute knockdown animals. Control and *ago-2* animals exhibit the expected lethality upon exposure to *histoneH3* dsRNA, while combined *ago-1+3* knockdown animals survive, consistent with impaired RNAi. Experiments were initiated after two weeks of argonaute knockdown.
**G.** Expression of *dcr-1* and *dcr-2* and neoblast markers (*bruli*, *vasa*, *mcm2*) following dicer knockdown, indicating preserved neoblast abundance.
**H.** Representative phenotypes following dicer knockdown. *dcr-2* RNAi results in head regression, whereas *dcr-1* RNAi and controls are phenotypically normal.
**I.** Survival analysis following dicer knockdown. *dcr-2* RNAi is lethal, whereas *dcr-1* RNAi and controls maintain viability.

#### Supplemental Figure S3

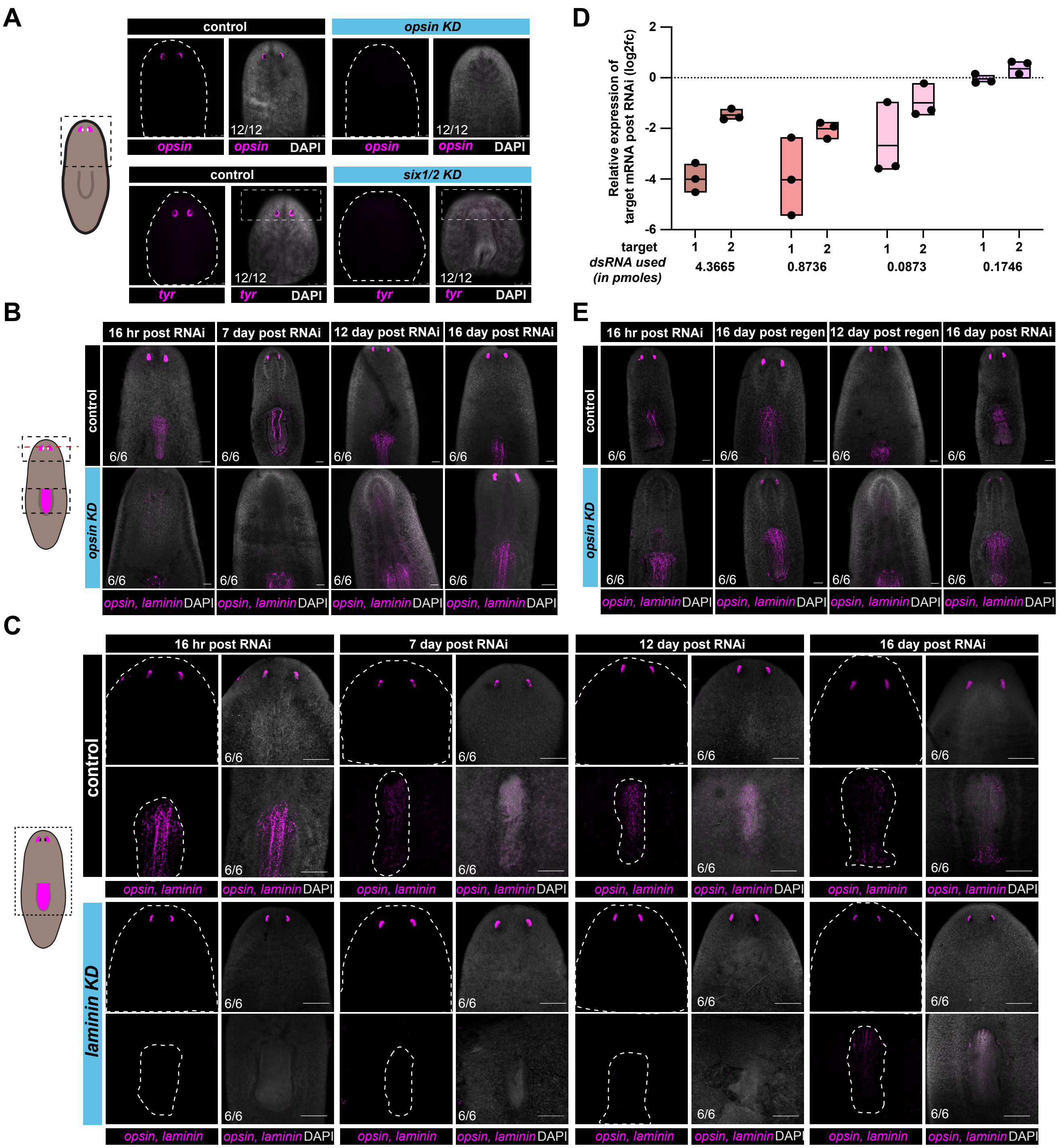

**Supplementary Figure S3. RNAi is specific and reversible.**
**A.** Confocal FISH images showing loss of eye-specific *opsin* mRNA (magenta) following *opsin* RNAi and loss of *tyrosinase* mRNA (magenta) following *six-1/2* RNAi after a single dsRNA injection, compared with control RNAi.
**B.** Confocal FISH images showing the specificity and reversibility of *opsin* RNAi. *Opsin* mRNA (magenta) progressively reappears over time (16 h, 7 d, 12 d, 16 d) following RNAi, while the non-target pharyngeal transcript *laminin* (magenta) remains unchanged. Some images are repeated from Main Figure 2B to show the relation of FISH signal to the outline of the animal shown by DAPI staining.
**C.** Confocal FISH images showing the specificity and reversibility of *laminin* RNAi. *Laminin* mRNA (magenta) reappears over time (16 h, 7 d, 12 d, 16 d) following RNAi, whereas the non-target eye-specific transcript *opsin* (magenta) is unaffected.
**D.** Box plots showing relative mRNA expression levels for two independent dsRNA targets across increasing dsRNA concentrations (pmol), as measured by qPCR.
**E.** Confocal FISH images showing recovery of *opsin* mRNA (magenta) during regeneration following RNAi (16 h, 12 d, 16 d), compared with homeostatic controls (16 d), while *laminin* expression remains unaffected. Some images are repeated from Main Figure 2C to show the relation of FISH signal to the outline of the animal shown by DAPI staining.

### Supplemental Figure S4

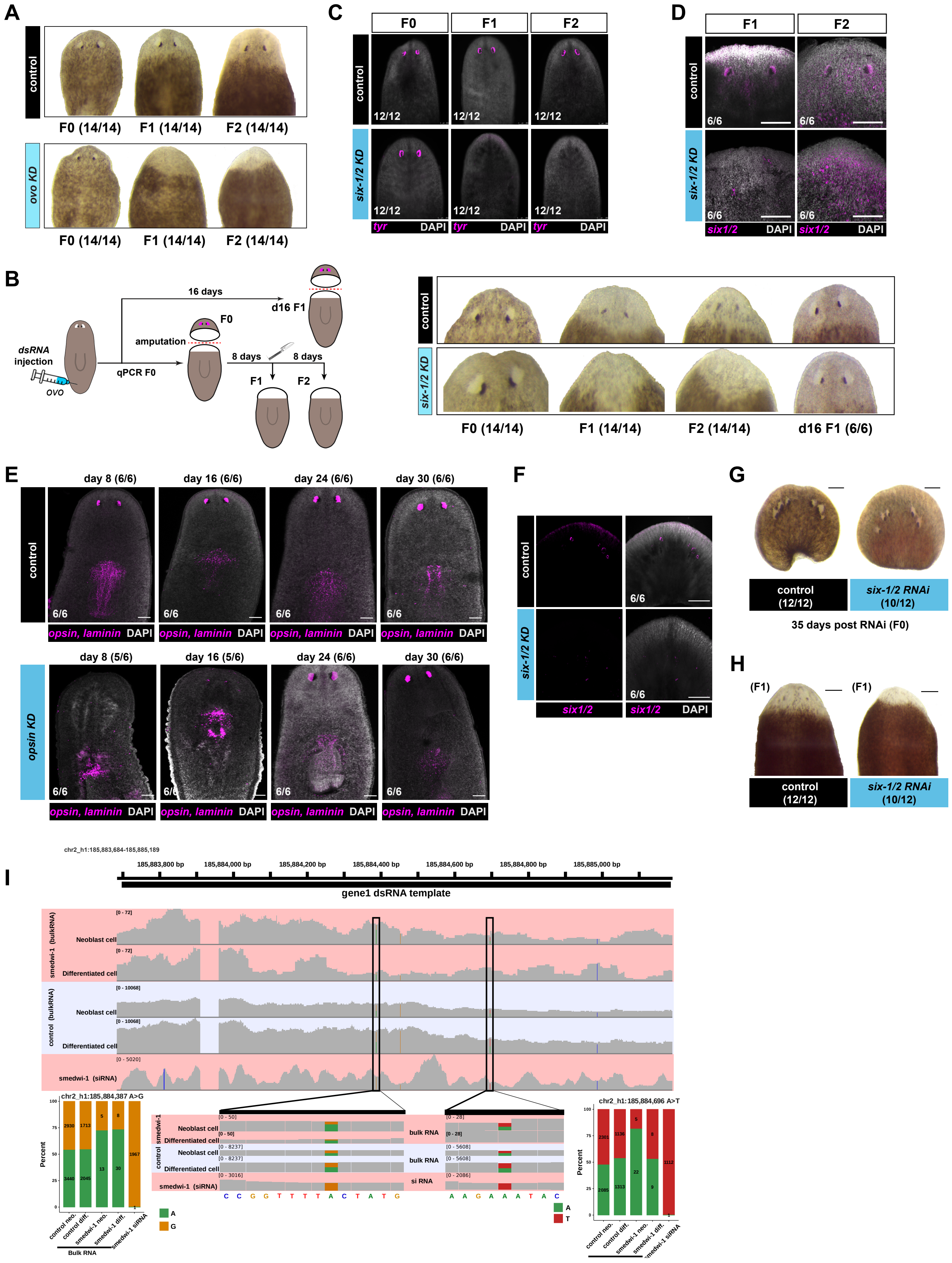

**Supplementary Figure S4. Phenotypic effects of RNAi across regeneration are not due to sustained target mRNA depletion.**
**A.** Bright-field images showing macroscopic eye phenotypes following *ovo* or control dsRNA injection combined with repeated head amputation and regeneration. Eye defects persist across multiple regeneration cycles. Some images are repeated from Main Figure 2D to show the comparison to control RNAi.
**B.** Experimental schematic (left) and bright-field images (right) showing macroscopic eye phenotypes following *six-1/2* or control dsRNA injection and repeated head amputation and regeneration, with defects maintained over successive regeneration cycles if amputatoins start early after injection. Amputation at 16d post injection however results in normal regenerated eyes.
**C.** Confocal FISH images showing loss of *tyrosinase* (*tyr*; magenta), a marker of differentiated eye cells, confirming sustained impairment of eye differentiation across multiple regeneration rounds.
**D.** Confocal FISH images showing re-expression of *six-1/2* mRNA (magenta) in the F2 generation following *six-1/2* RNAi, indicating that long-term phenotypic effects are not due to persistent depletion of the target transcript.
**E.** Confocal FISH images showing prolonged suppression of *opsin* mRNA (magenta) following dsRNA feeding, with RNAi effects persisting for at least one additional week as compared to dsRNA induced by injection (compare to Main Figure 2B).
**F.** Confocal FISH images showing reduction of *six-1/2* mRNA (magenta) in *Schmidtea polychroa* following *six-1/2* RNAi.
**G.** Brightfield images showing shrinking of macroscopic eyes in homeostatic heads in *Schmidtea polychroa* following *six-1/2* RNAi.
**H.** Brightfield images showing loss of macroscopic eyes in the regenerated heads of *Schmidtea polychroa* following *six-1/2* RNAi.
**I.** Genome browser tracks showing read coverage from bulk RNA-seq (control and *smedwi-1* RNAi) and small RNA-seq (*smedwi-1* RNAi) across *smedwi* genes and the *smedwi* dsRNA template. Read coverage is shown in gray, with putative single-nucleotide polymorphisms (SNPs) highlighted in color. Two SNPs identified in bulk RNA-seq-chr2_h1:185,884,387 (A>G; left) and chr2_h1:185,884,696 (A>T; right), are shown in detail, with allele frequencies compared across the indicated datasets.

#### Supplemental Figure S5

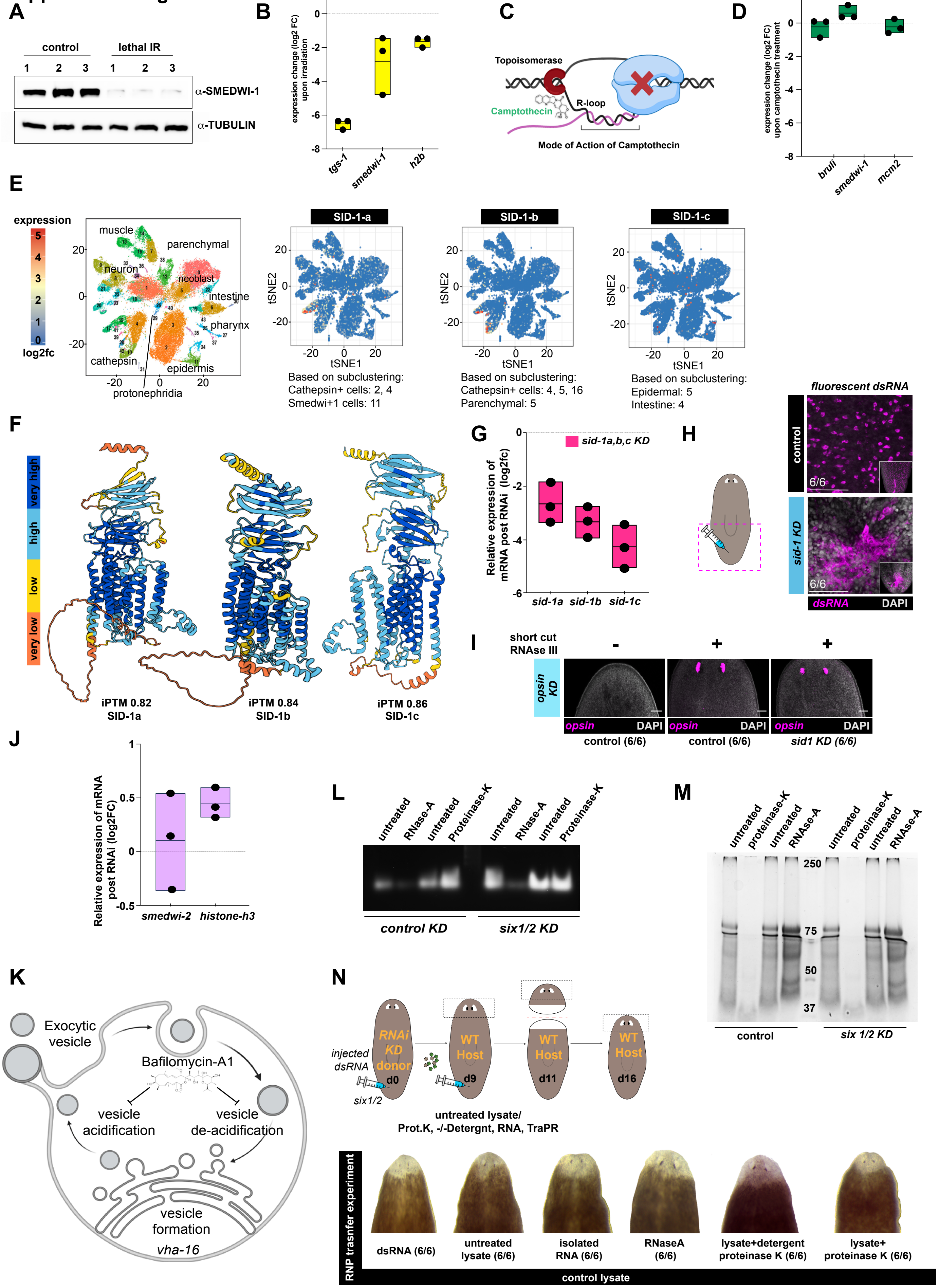

**Supplementary Figure S5. Stem cells initiate and RNPs propagate systemic RNAi in planarians.**
**A.** Western blot showing loss of the neoblast marker SMEDWI-1 six days after lethal irradiation (3000 rad), confirming ablation of stem cells.
**B.** Box plots showing reduction in the mRNA levels of neoblasts genes *tgs-1, smedwi-1,* and *histoneH2b* upon irradiation, as quantified by qPCR.
**C.** Model illustrating the mode of action of camptothecin, which inhibits DNA topoisomerase I (Top1), causing replication arrest and S-phase accumulation (Pommier 2006).
**D.** Box plots showing minimal changes in the mRNA levels of neoblasts genes *bruli-1, smedwi-1,* and *mcm2* upon Camptothecin treatment, as quantified by qPCR.
**E.** UMAP visualization of single-cell RNA-seq data (Fincher et al. 2018) showing planarian tissue types (left) and expression patterns of *sid-1* homologues (right).
**F.** AlphaFold-predicted models of *S. mediterranea* SID-1 homologues, colored by per-residue confidence (pLDDT), with high-confidence regions (blue; pLDDT > 90) and low-confidence regions (orange; pLDDT < 50).
**G.** Box plots showing reduced expression of *sid-1* homologues following combined knockdown, as determined by qPCR.
**H.** Experimental schematic (left) and confocal FISH images showing localization of injected dsRNA in control animals and *sid*-1 knockdown animals. In control animals, dsRNA is imported into cells, whereas this is impaired following *sid-1* knockdown.
**I.** Confocal FISH images showing *opsin* mRNA after injection of *opsin* dsRNA with or without RNase III pretreatment in control and *sid-1* knockdown animals. RNase III–treated dsRNA fails to induce RNAi.
**J.** Box plots showing ineffective knockdown of *smedwi-2* and *histone H3* following test RNAi in *sid-1* knockdown animals, normalized to combined *sid-1* knockdown with control dsRNA injection.
**K.** Model illustrating the proposed roles of VHA-16 and Bafilomycin A1 (Bayer et al. 1998) in endocytic and exocytic pathways.
**L.** Agarose gel stained with SYBR Gold, showing total RNA integrity in the indicated lysates.
**M.** Polyacrylamide gel stained with PAGE-Blue showing total protein from the indicated lysates.
**N.** Schematic (top) of the RNAi transfer assay in which lysates from control animals subjected to the indicated treatments were injected into wild-type hosts. Bright-field images (bottom) show no loss of eye structures following transfer of either treated or untreated lysates.

**Supplemental Table 1. Primer sequences used in this study.**

**Supplemental Table 2. IDs of proteins used in phylogenetic analysis.**

**Table S1: qPCR Primers used in this study**

| name | sequence |
|---|---|
| agat-1F | CCTAAAAGGCGAAGGTGTGACT |
| agat-1R | TGCAACATCCAAACCGACAGA |
| agat-3F | ACTGCTAAGTGAAGAACGACTAT |
| agat-3R | TAAACACTCGACCCGGATCC |
| ago-1F | GAAATTGCCGCCATTCAATATGC |
| ago-1R | ACCGAATCCTCTAAAGTAATTGGAC |
| ago-2F | CATATGTCAGATGTACAAGATCCGT |
| ago-2R | GGGAGCCTTCTCCACTATCG |
| ago-3F | TTCCTCAAAATGAAAAAGATGGAGT |
| ago-3R | CCTTGTATGCCTTCGTGTGA |
| bruliF | ATGATATTACAGCAATTTCCACATGCCA |
| bruliR | GCGGAGGTGTTAAATTTGGGCT |
| collagenF | TGGTGTTCCAGGAAGAGAAGGT |
| collagenR | GTGGCCCTGGATCACCTTCA |
| dicer-1F | CCAGTTCCATCTGCAGCGAA |
| dicer-1R | TGCCGCCTTATGCCAAAATGT |
| dicer-2F | GCGCACGAGCTCTACCTTTA |
| dicer2-R | TGTGATTGAAGAAGGTCTTGACGT |
| gapdhF | TCTTCCCCAACCAATTTTCTGTTCTG |
| gapdhR | CCGAATATTTTATTTGGCTCTTCCTCCA |
| histone-h2bF | TAACCAGCAGAGAGATTCAAACTG |
| histone-h2bR | GTTCCTTCACTAACTGCGTGTTTA |
| histone-h3F | GCGATTTCTCGCACCAGACG |
| histone-h3F | CACCTGCAACTGGTGGTGTTAAG |
| mcm2F | TAACATCAATTTCATCTCCAGGCTTGC |
| mcm2R | GAGAGTCCCGGTACTGTTCCG |
| prog-1F | GCAATCTGCTTTCGTAATGTGTCCT |
| prog-1R | TCTGCAAAGTCTCCCGCCAA |
| sid1aF | GATGATGCCGGCGATTGCTC |
| sid1aR | ATAGCAATGGTTTGTTGGGGGT |
| sid1bF | GGATTGGCTCAATAACTACCGCAT |
| sid1bR | CCACAAGCTCCAGTGACATCG |
| sid1cF | ACAAAGGAAGCTGCATTAAGGGT |
| sid1cR | GCTGCCTTTACCGGTACTTCG |
| smedwi-1F | GTCTCAGAAAACAACTAAAGGTACAGCA |
| smedwi-1R | TGCTGCAATACACTCGGAGACA |
| smedwi-2F | ATGGAGGAAATACCAGTAAAAGTAGCT |
| smedwi-2R | TTCTAAATGTTCTTCTCAGGCCACC |
| tgs-1F | GCCAACACTGAAAAATCAAACGTTGA |
| tgs-1R | CGCTGCTAGCTTTCGCTTCT |
| ubiquilinF | AAATTCGCCTGCCTGTTGGG |
| ubiquilinR | CCGGTGGCATTAATCCATCTGT |
| vasaF | CGACTGCCGATGCTGGAGAT |
| vasaR | CCTCATGGCATGCGCACAAA |
| vimentinF | AACCGCGGCTTCAACTGAAC |
| vimentinR | CAGCGGAACTTAAAACTCGCTCTT |
| zfp-1F | CCCGTGCCTGAACAATTTGACA |
| zfp-1R | CCTCAGCGCATGCCTCTGTA |
| ovo_RT | TTTTTTTTTTTTTTTTTTAAATTTGTTTTTAATTATTCGATATAAAA |
| ovo-F | ACAACTGAGCATCGAAGTTAATCACA |
| ovo-R | ACACCGTGTGAACGTATATGACG |
